# The blast pathogen effector AVR-Pik binds and stabilizes rice heavy metal-associated (HMA) proteins to co-opt their function in immunity

**DOI:** 10.1101/2020.12.01.406389

**Authors:** Kaori Oikawa, Koki Fujisaki, Motoki Shimizu, Takumi Takeda, Keiichiro Nemoto, Hiromasa Saitoh, Akiko Hirabuchi, Yukie Hiraka, Naomi Miyaji, Aleksandra Białas, Thorsten Langner, Ronny Kellner, Tolga O Bozkurt, Stella Cesari, Thomas Kroj, Mark J Banfield, Sophien Kamoun, Ryohei Terauchi

## Abstract

Intracellular nucleotide-binding domain and leucine-rich repeat-containing (NLR) receptors play crucial roles in immunity across multiple domains of life. In plants, a subset of NLRs contain noncanonical integrated domains that are thought to have evolved from host targets of pathogen effectors to serve as pathogen baits. However, the functions of host proteins with similarity to NLR integrated domains and the extent to which they are targeted by pathogen effectors remain largely unknown. Here, we show that the blast fungus effector AVR-Pik binds a subset of related rice proteins containing a heavy metal-associated (HMA) domain, one of the domains that has repeatedly integrated into plant NLR immune receptors. We find that AVR-Pik binding stabilizes the rice small HMA (sHMA) proteins OsHIPP19 and OsHIPP20. Knockout of *OsHIPP20* causes enhanced disease resistance towards the blast pathogen, indicating that *OsHIPP20* is a susceptibility gene (*S*-gene). We propose that AVR-Pik has evolved to bind HMA domain proteins and co-opt their function to suppress immunity. Yet this binding carries a trade-off, it triggers immunity in plants carrying NLR receptors with integrated HMA domains.

**Significance Statement:** Rice blast disease, caused by the fungus *Magnaporthe oryzae*, is one of the most devastating diseases of rice. Therefore, understanding the mechanisms of blast fungus infection and resistance of rice against the disease is important for global food security. In this study, we show that the *M. oryzae* effector protein AVR-PikD binds rice sHMA proteins and stabilizes them, presumably to enhance pathogen infection. We show that loss-of-function mutants in one rice sHMA, OsHIPP20, reduced the level of susceptibility against a compatible isolate of *M. oryzae*, suggesting that *M. oryzae* requires host sHMA to facilitate invasion. Remarkably, *OsHIPP20* knockout rice line showed no growth defect, suggesting editing sHMA genes may present a novel source of resistance against blast disease.

## Introduction

Plant pathogens target host processes to promote disease by secreting effector proteins (^1^Hogenhout et al. 2009). Some of the host targets of effectors have been co-opted by plant intracellular nucleotide-binding leucine rich repeat (NLR) immune receptors to act as baits to detect pathogens, and in this context are known as integrated domains (IDs) (^2^Cesari et al. 2014; ^3^Wu et al. 2015; ^4^Sarris et al. 2016; ^5^Kroj et al. 2016). Genome-wide bioinformatics searches have found such domains in diverse NLR immune receptors from multiple plant families (^4^Sarris et al. 2016; ^5^Kroj et al. 2016; ^6^Baggs et al. 2017; ^7^Bailey et al. 2018). We hypothesize that such widespread NLR-IDs modulate basic immune responses that are conserved among plants. One such domain is the heavy metal-associated (HMA) domain found in four botanical families (^4^Sarris et al. 2016). In rice, HMA domains have been integrated into two different NLRs, RGA5 (^8^Okuyama et al. 2011) and Pik-1 (^9^Ashikawa et al. 2008). In addition to rice (a member of Poaceae), HMA domains have also integrated into NLR immune receptors of plant species in the Brassicaceae, Fabaceae and Rosaceae (^4^Sarris et al. 2016; ^5^Kroj et al. 2016). This indicates that HMA-containing proteins have likely been repeatedly targeted by pathogens across the diversity of flowering plants. Therefore, understanding the endogenous function of HMA-containing proteins has the potential to reveal important basic features of plant disease susceptibility and immunity. In this study, we report rice HMA-containing proteins that are targets of a blast pathogen effector from *Magnaporthe* (syn. *Pyricularia*) *oryzae* and address their potential function.

The rice *Pik* locus comprises two NLR genes, *Pik-1* and *Pik-2*, and recognizes the *M. oryzae* effector AVR-Pik (^9^Ashikawa et al. 2008), triggering an immune response that restricts infection. AVR-Pik is a 113-amino-acid protein originally defined as having no sequence similarity to known protein domains (^10^Yoshida et al. 2009). More recently, structure-informed similarity searches showed that AVR-Pik belongs to the MAX (*Magnaporthe* AVRs and ToxB-like) family of fungal effectors (^11^De Guillen et al. 2015), which adopt a six β-sandwich fold stabilized by buried hydrophobic residues, and commonly but not always, a disulphide bond. Pik-1 recognition of AVR-Pik is mediated by direct binding of the effector to an HMA domain (^12^Maqbool et al. 2015) located between the N-terminal coiled-coil (CC) and nucleotide binding (NB) domains of Pik-1 (^13^Kanzaki et al. 2012; ^14^Cesari et al. 2013). *AVR-Pik* and *Pik-1* are described as being involved in a coevolutionary arms race that has resulted in the emergence of allelic series of both effector genes in the pathogen and NLR genes in the host (^13^Kanzaki et al. 2012; ^15^De la Concepcion et al. 2018; ^16^Białas et al. 2018).

Biochemical and structural analysis of complexes between AVR-Pik variants and HMA domains of different Pik-1 alleles revealed the molecular interactions between the effector and NLR-ID (^12^Maqbool et al. 2015; ^15^De la Concepcion et al. 2018; ^17^De la Concepcion et al. 2019). This knowledge recently allowed structure-guided protein engineering to expand the recognition profile of a Pik NLR to different AVR-Pik variants (^17^De la Concepcion et al. 2019; ^18^Maidment et al. 2023). The Pik-1 HMA domains exhibit a four β-sheets and two α-strands (βαββαβ) topology similar to the yeast copper transporter domain Ccc2A (^19^Banci et al. 2001), even though the characteristic MxCxxC metal-binding motif is degenerate in Pik-1. The integrated HMA domain of RGA5 also adopts the classical HMA domain fold but, intriguingly, uses a different interface to interact with the *M. oryzae* effectors AVR-Pia and AVR1-CO39 (^20^Guo et al. 2018).

HMA domains are also found in other plant proteins that are unrelated to NLRs (^21^De Abreu-Neto et al. 2013). These proteins form large and complex families known as heavy metal-associated plant proteins (HPPs) and heavy metal-associated isoprenylated plant proteins (HIPPs), here collectively referred to as small proteins containing an HMA domain (abbreviated as sHMA proteins). One such sHMA protein is the product of the rice blast partial resistance gene *pi21* (^22^Fukuoka et al. 2009). The recessive allele *pi21*, a presumed loss-of-function allele with a deletion mutation, confers partial broad-spectrum resistance to rice against compatible isolates of *M. oryzae.* This finding implicates HMA domain-containing proteins in rice defense (^22^Fukuoka et al. 2009). However, the molecular function of Pi21 and other rice sHMA proteins have not been characterized to date.

Unlike other *M. oryzae* effectors, *AVR-Pik* does not show extensive presence/absence polymorphisms within the rice-infecting lineage, and its evolution in natural pathogen populations is mainly driven by nonsynonymous amino acid substitutions (^13^Kanzaki et al. 2012; ^23^Shi et al. 2018). This suggests that *AVR-Pik* encodes an activity of benefit to the pathogen that is maintained in resistance-evading forms of the effector. To address the virulence function of AVR-Pik, we set out to identify rice proteins other than the Pik NLRs that interact with this effector. We found that AVR-Pik binds and stabilizes a subset of sHMA proteins. Knockout of one sHMA gene (*OsHIPP20*) conferred enhanced resistance to infection by the blast pathogen, suggesting *OsHIPP20* is a susceptibility gene (S-gene). Our model is that AVR-Pik effectors interfere with sHMA function by stabilizing and relocating these proteins to support pathogen invasion.

## Results

### AVR-PikD binds members of a subclade of small heavy metal associated proteins (sHMAs) of rice

To identify rice proteins that may be putative targets of AVR-PikD, we performed a yeast 2-hybrid screen (Y2H) with the effector as bait and a cDNA library prepared from leaves of rice cultivar Sasanishiki inoculated with *M. oryzae* as the prey. From this screen we identified four HMA-containing proteins, named OsHIPP19 (LOC_Os04g39350), OsHIPP20 (LOC_Os04g39010), OsHPP04 (LOC_Os02g37300) and OsHPP03 (LOC_Os02g37290) (^21^De Abreu-Neto et al. 2013), as interactors of AVR-PikD, amongst other proteins (**Table S1**). The sizes of AVR-PikD interacting HMAs ranged from 118 (OsHPP03) to 123 (OsHIPP19) amino acids.

Rice sHMA proteins typically comprise a conserved N-terminal HMA domain followed by a variable proline-rich domain (**Fig. 1A**) and may contain a C-terminal “CaaX” isoprenylation motif (where “a” represents an aliphatic amino acid and X represents any amino acid). They form a large protein family with 87 members in the rice genome (cultivar Nipponbare) as annotated by Rice Genome Annotation Project (^24^Kawahara et al. 2013) (**Fig. 1B**). Phylogenetic analyses of the aligned HMA domains of rice sHMA proteins revealed two clades supported by high bootstrap values (> 90%) that we designate here as Clades A and B. All four sHMA proteins interacting with AVR-PikD belong to Clade A (**Fig. 1B**). Interestingly, the HMA domains of RGA5 and three alleles of the Pik-1 NLRs also cluster in Clade A. However, the integrated HMA domains of Pik-1 (Pik*-HMA, Pikm-HMA and Pikp-HMA) and RGA5 (RGA5-HMA) are on separate branches in the tree, indicating distinct lineages and diversification patterns.

**Fig. 1.**
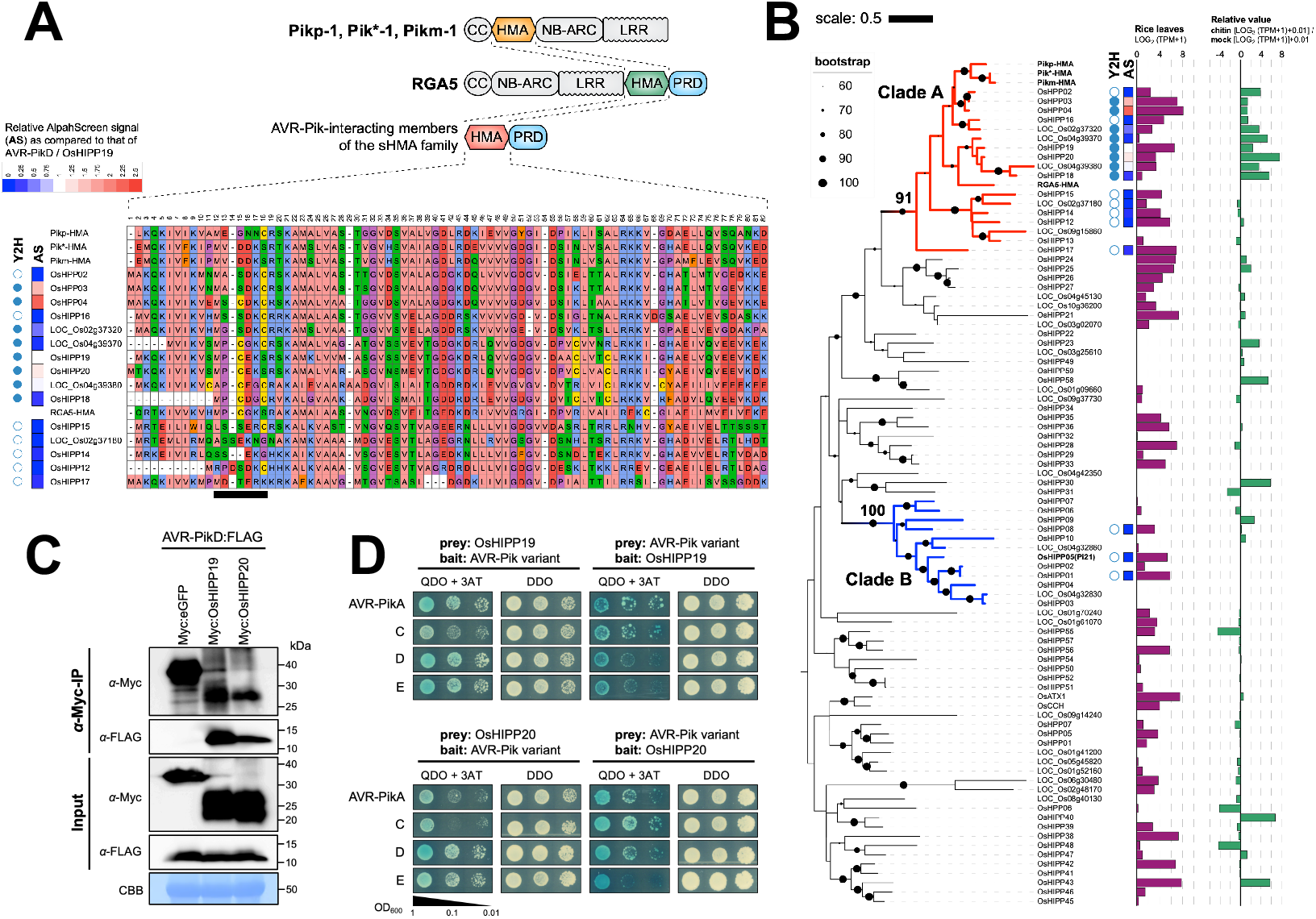
AVR-PikD binds Clade A sHMAs. **A:** Schematic representation of the Pik-1 (Pikp-1, Pik*-1, Pikm-1) and RGA5 Nucleotide-binding Leucine Rich Repeat Receptors (NLRs) and small HMA (sHMA) proteins of rice. CC: coiled-coil domain; NB-ARC: nucleotide binding domain; LRR: leucine rich repeat; PRD: proline-rich domain. Amino acid sequence alignment of a subset of HMA proteins of rice. The black bar highlights the putative metal-binding motif MxCxxC. Interaction with AVR-PikD is indicated for yeast two-hybrid (Y2H: blue dot: binding; white dot: non-binding) and AlphaScreen (AlphaScreen Signal [AS]: strength of interaction signal as compared to that of AVR-PikD/OsHIPP19 interaction is given in the inset). **B:** A maximum likelihood tree of the HMA domains of 87 sHMA of rice. Amino acid sequences of the HMA domains were aligned and used for reconstruction of the phylogenetic tree. The dots on the branches indicate bootstrap values after 1,000 replications. Clade A and Clade B are indicated by red and blue branches, respectively. Note Pi21 (OsHIPP05) belongs to Clade B. Y2H and AlphaScreen results (AS) are as shown as in A. Purple bar graphs (log2(TPM+1) value) show expression level of each gene in leaves as revealed by RNA-seq. Green bar graphs show induction levels of each gene in rice suspension cultured cells after chitin treatment (log2[TPM of chitin-treated cultured cells+1] + 0.01)/ (log2[TPM of mock-treated cultured cells+1] + 0.01). **C:** Results of co-immunoprecipitation of AVR-PikD with OsHIPP19 and OsHIPP20 transiently expressed in Nicotiana benthamiana leaves. **D:** Y2H interactions between variants of AVR-Pik; AVR-PikA, AVR-PikC, AVR-PikD, AVR-PikE against OsHIPP19 and OsHIPP20.

To determine if AVR-PikD interacts with other Clade A sHMA proteins, we selected 15 that are expressed in rice leaves (**Fig. 1B**) and tested pairwise interactions by Y2H (**Fig. S1**). This experiment showed that AVR-PikD binds around half of the tested Clade A sHMA proteins (**Fig. 1B**). We also tested binding of AVR-PikD with three sHMA proteins from Clade B, including OsHIPP05 (Pi21). These sHMA proteins did not bind AVR-PikD (**Fig S1**), revealing that AVR-PikD shows specific binding to Clade A sHMA proteins (**Fig. 1B**). These results were overall confirmed using the AlphaScreen (Amplified Luminescent Proximity Homogenous Assay Screen) method (^25^Ullman et al. 1996; ^26^Nemoto et al. 2017) (**Fig. 1B**; **Fig. S2**). Interestingly, RNA-seq analysis indicated that AVR-PikD-interacting sHMAs were induced by chitin treatment of rice suspension cultured cells (**Fig. 1B**), suggesting potential roles in immunity.

We also tested the interaction of Clade A sHMA proteins OsHIPP19, OsHIPP20, OsHPP04, OsHPP03 and LOC_Os04g39380 with the *M. oryzae* effectors AVR-Pia and AVR1-CO39, which interact with the HMA domain of RGA5 (^14^Cesari et al. 2013). We found that AVR-Pia and AVR1-CO39 did not bind any of the sHMAs tested (**Fig. S3**). Also, none of the three AVRs interacted with the Pi21 HMA protein (**Fig S3**). Interactions between AVR-PikD and OsHIPP19 and OsHIPP20 were further confirmed by co-immunoprecipitation using proteins expressed in *N. benthamiana* (**Fig. 1C**). Naturally occurring AVR-Pik variants are differentially recognized by allelic Pik NLRs. These recognition specificities correlate with the binding affinity of AVR-Pik variants to the integrated HMA domain of the Pik-1 NLR (^13^Kanzaki et al. 2012; ^12^Maqbool et al. 2015; ^15,16^De la Concepcion et al. 2018; 2019). We tested whether the AVR-Pik variants AVR-PikA, C, or E, interact with the rice sHMA proteins OsHIPP19 and OsHIPP20 in Y2H. The results of this experiment (**Fig. 1D; Fig. S4**) showed that similar to AVR-PikD, all AVR-Pik variants tested interacted with OsHIPP19 and OsHIPP20. This result suggests that all the tested AVR-Pik variants bind sHMAs, the possible host target proteins, whereas they vary in the recognition by different alleles of Pik NLRs.

After posting the first version of this preprint to bioRxiv in 2020, ^27^Maidment (2021) applied gel filtration assay and Surface Plasmon Resonance (SPR) assay and confirmed that OsHIPP19 binds AVR-PikD, C and F with nanomolar affinity.

### Three regions of sHMAs have close contact with AVR-PikD and model-guided structure prediction allowed converting AVR-Pik non-binding sHMAs to AVR-Pik-binding

A previous study by ^27^Maidment et al. (2021) determined the crystal structure of the AVR-PikF/OsHIPP19 complex, which was similar to that of AVR-PikD/Pikp1-HMA (^12^Maqbool et al. 2015) and described three predominant interfaces. We took advantage of the multiple interactions we identified in our Y2H screen to further investigate AVR-PikD/sHMA complexes using computational structural biology. We predicted AVR-PikD/sHMA complex models using ColabFold v1.5.2 (AlphaFold2 using MMseqs2) (^28^Mirdita et al. 2022), which revealed that three regions A, B, C conserved in Clade A sHMA seem closely (Atomic distance < 5Å) positioned to AVR-PikD (**Fig. 2A**; **Fig. S5**). These regions correspond to the interfaces 1, 2, 3, respectively, as reported by ^15^De la Concepcion et al. (2018) and ^27^Maidment et al. (2021). To validate the importance of these interfaces, we selected OsHIPP16, an sHMA in Clade A that does not bind AVR-PikD in Y2H (**Fig. S1**) or AlphaScreen (**Fig. S2**), for further investigation. Modeling of the interaction predicted that OsHIPP16 does not form a complex with AVR-PikD in the way experimentally determined for OsHIPP19/AVR-PikF or predicted for OsHIPP20/AVR-PikD (**Fig. 2A; Fig. S5C**). We compared the amino acid sequences of OsHIPP16 and OsHIPP20 and found that the former has an insertion of an aspartic acid (D) at position 68, and has a serine (S) at position 80 where a lysine (K) is present in OsHIPP20 (**Fig. 2A**). Modeling indicated a deletion of D68 (resulting in OsHIPP16-D68DEL) and an amino acid replacement S80K (OsHIPP16-S80K) could render OsHIPP16 capable of binding AVR-PikD (**Fig. 2B; Fig. S6**). In a Y2H assay, OsHIPP16-D68DEL did not bind AVR-PikD, but OsHIPP16-S80K bound the effector, and OsHIPP16-D68DEL/S80K strongly bound (**Fig. 2C; Fig. S7**). In an AlphaScreen experiment, each single mutant weakly bound AVR-PikD, but OsHIPP16-D68DEL/S80K strongly bound AVR-PikD (**Fig. 2D).** We also tested a second AVR-PikD non-binding sHMA, OsHPP02 (**Fig. S1, S2**) to see whether we could obtain a gain-of-binding mutant for this protein. Modeling of the interaction predicted that OsHPP02 does not form an OsHIPP20/AVR-PikD-like complex (**Fig. S5C**), but may be converted to AVR-PikD-binding by a single amino acid change at position 79 (within region C) from an aspartic acid (D) to the valine (V) found in OsHIPP20 (**Fig. 2A**; **Fig. S8**). Indeed, OsHPP02-D79V bound to AVR-PikD in both Y2H and AlphaScreen assays (**Fig. S8**). Overall, these results, obtained by model-based interaction prediction, confirm the findings previously observed using the AVR-PikF/OsHIPP19 crystal structure (^27^Maidment et al. 2021) that region C conserved in Clade A sHMAs is important for the interactions between AVR-PikD and Clade A sHMAs.

**Fig. 2.**
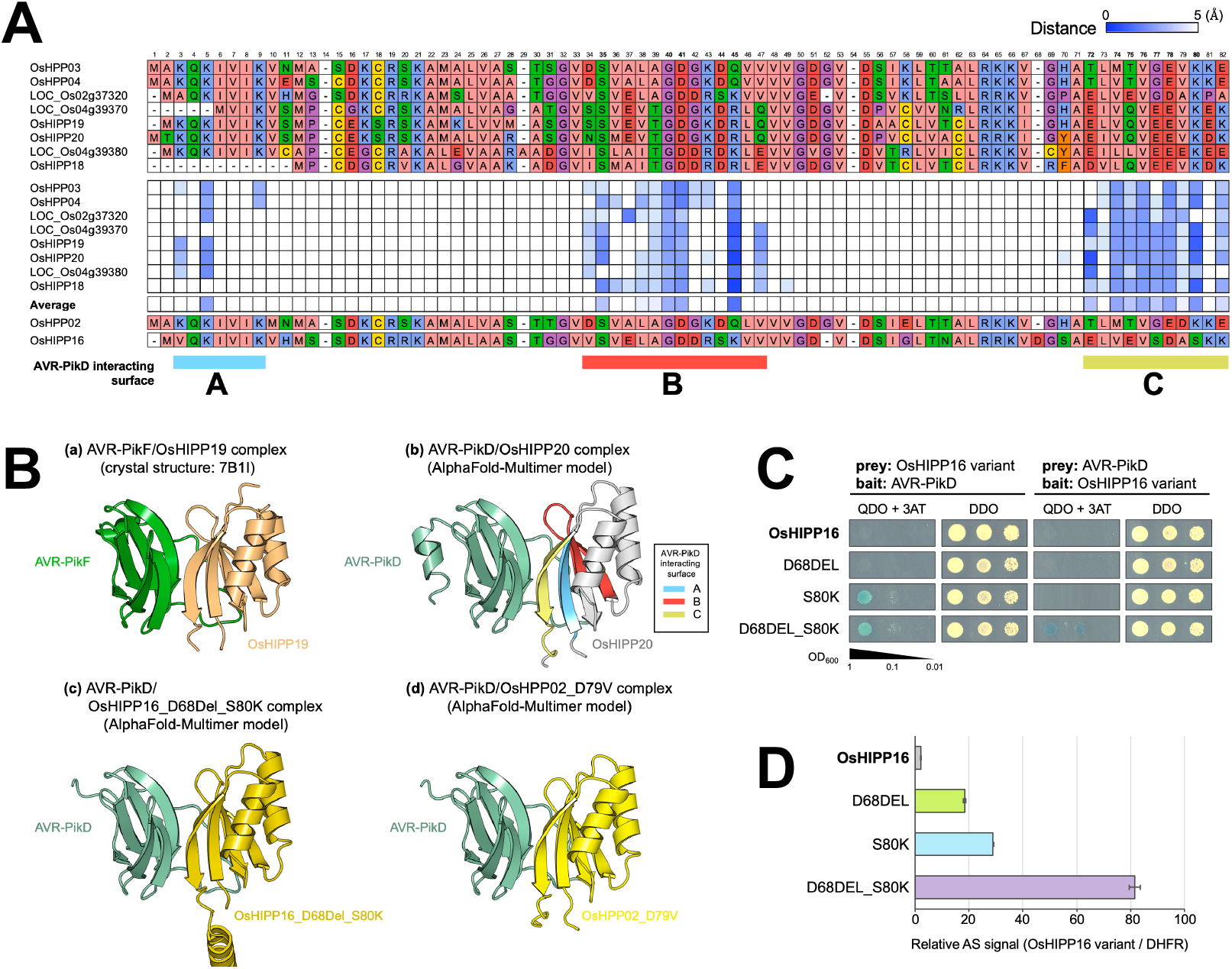
Three regions of sHMAs are predicted to have close contact with AVR-PikD. **A:** Amino acid sequence alignment of sHMAs (top) and a matrix of distance of each amino acid residue to AVR-PikD protein as predicted by ColabFold. Predicted atomic distance below 5Å are indicated by blue tiles. Three regions indicated by A, B and C seem in close contact with AVR-PikD. **B:** Binding structures between AVR-PikF (green) and OsHIPP19 (light orange) (a: Crystal structure; Maidment et al. 2021), predicted structure between AVR-PikD (light green) and OsHIPP20 (grey) (b: AlphaFold-multimer model), predicted structure between AVR-PikD (light green) and OsHIPP16_D68DELS80K (dark yellow) (c: AlphaFold-multimer model), and predicted structure between AVR-PikD (light green) and OsHPP02_D79V (yellow) (d: AlphaFold-multimer model). **C:** Y2H interactions between the variants of OsHIPP16 (OsHIPP16, OsHIPP16_D68DEL, OsHIPP16_S80K, OsHIPP16_D68DEL_S80K) and AVR-PikD. **D:** AlphaScreen interactions between the variants of OsHIPP16 (OsHIPP16, OsHIPP16_D68DEL, OsHIPP16_S80K,OsHIPP16_D68DEL_S80K) and AVR-PikD. The values are relative AlphaScreen signals (AS) to that of OsHIPP16/DHFR interaction signal (negative control).

### AVR-PikD stabilizes OsHIPP19 and OsHIPP20 and affects their subcellular localization

Next, we aimed to determine the effect of AVR-PikD binding on the putative function sHMA proteins. Firstly, we co-expressed OsHIPP19 and OsHIPP20 with AVR-PikD in *N. benthamiana* leaves by agroinfiltration (see Materials and Methods). Following expression, leaf extract was separated into supernatant and pellet fractions by centrifugation, and each fraction was analyzed by western blot (**Fig. 3A; Fig. S9**). We used expression of β-glucuronidase (GUS) protein and AVR-Pii, a *M. oryzae* effector unrelated to AVR-Pik, as controls that do not bind OsHIPP19 and OsHIPP20. In the supernatant fraction we observed that the OsHIPP19 and OsHIPP20 proteins were degraded to smaller fragments when co-expressed with GUS or AVR-Pii. However, when co-expressed with AVR-PikD, we detected stronger signals of intact OsHIPP19 and OsHIPP20. As an additional control we also tested OsHIPP17, an sHMA that does not bind AVR-PikD in either Y2H or by in planta coimmunoprecipitation, and only weakly interacts with AVR-PikD by the AlphaScreen method (**Fig. 1, 3A, Fig. S2; S10)**. We found that OsHIPP17 was degraded to a smaller fragment even in the presence of AVR-PikD. These results show that the AVR-PikD effector stabilizes sHMA proteins OsHIPP19 and OsHIPP20 in the plant cytosol, and this stabilization is specific to sHMA proteins that interact with the effector. We also noted consistently observed lower accumulation of AVR-PikD in the supernatant when co-expressed with OsHIPP17. This suggests the possibility that AVR-PikD may require interaction with other proteins in the plant cytosol for its own stability.

**Fig. 3.**
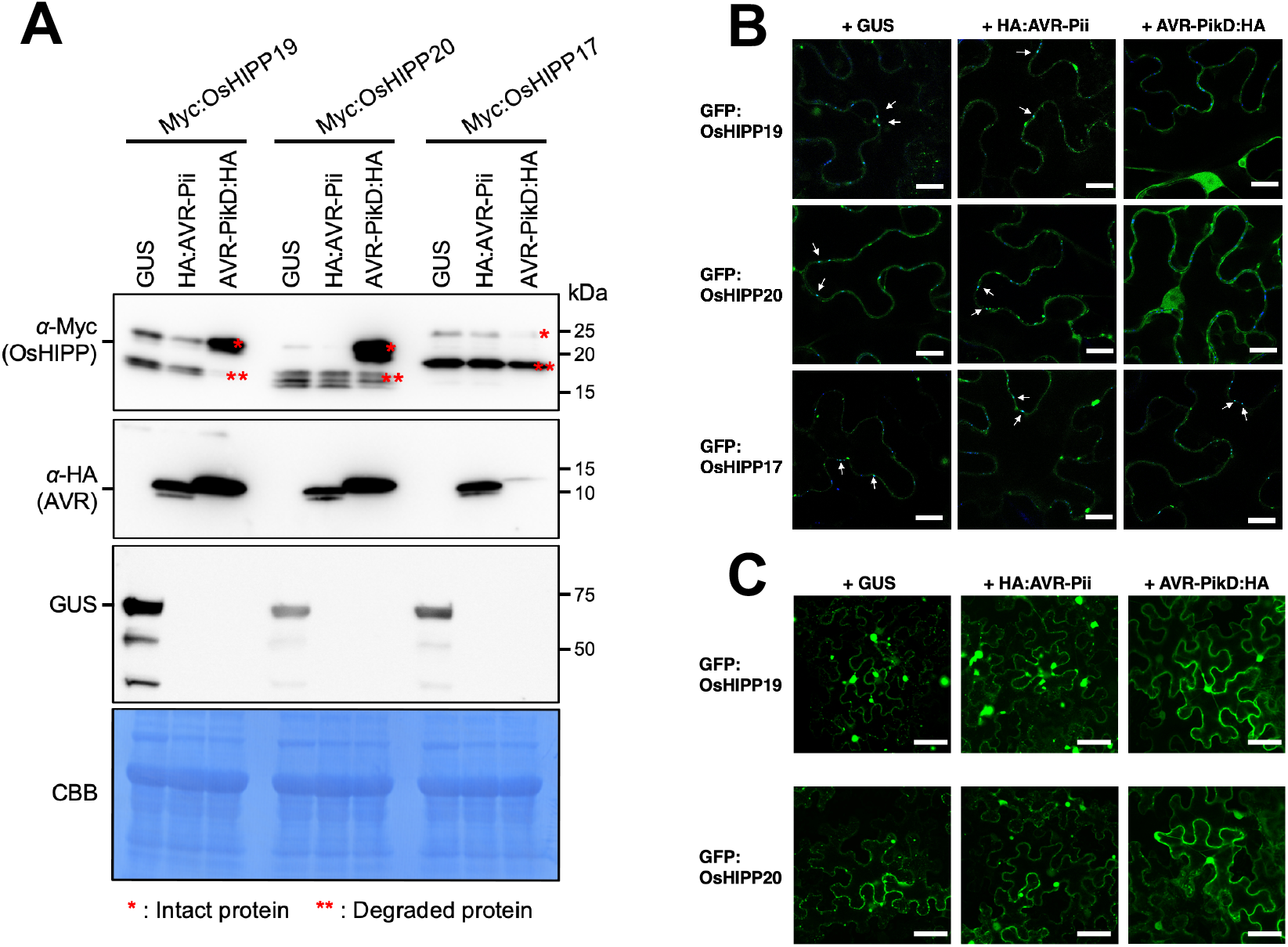
AVR-PikD stabilizes sHMA proteins and alters sHMA subcellular localization in *N. benthamiana*. **A:** sHMA proteins (Myc:OsHIPP19, Myc:OsHIPP20 and Myc:OsHIPP17) were transiently expressed in *N. benthamiana* leaves together with either GUS, HA:AVR-Pii or AVR-PikD:HA and were detected by an anti-Myc antibody. The result for supernatant fraction after fractionation of leaf extract is shown. The result for pellet fraction is shown in Fig. S9. OsHIPP19 and OsHIPP20 bound by AVR-PikD remain largely stable, whereas OsHIPP19 and OsHIPP20 expressed with GUS or AVR-Pii were degraded to a lower mass fragment. OsHIPP17 binds AVR-PikD only weakly and is degraded even in the presence of the effector. AVR-PikD seems unstable when unbound to target proteins. **B:** GFP:OsHIPP19 and GFP:OsHIPP20 seems to accumulate at plasmodesmata. Plasmodesmata are stained by aniline blue (blue color). White arrows indicate colocalization of GFP and aniline blue (Cyan color). Co-expression of AVR-PikD:HA relocates GFP:OsHIPP19 and GFP:OsHIPP20 from plasmodesmata. GFP:OsHIPP17 co-expressed with AVR-PikD:HA shows no relocation. Scale bar: 20 μm. **C:** GFP:OsHIPP19 and GFP:OsHIPP20 accumulate to punctae-like structures in the cells when expressed with GUS or HA:AVR-Pii, whereas these proteins were evenly distributed in the cytoplasm when expressed with AVR-PikD:HA. We obtained similar results in three independent experiments. Scale bar: 200 μm.

Next, we tested whether binding of AVR-PikD affected the subcellular localization of specific sHMA proteins. We transiently co-expressed GFP-tagged OsHIPP19 and OsHIPP20 (GFP:OsHIPP19 and GFP:OsHIPP20) by agroinfiltration in *N. benthamiana* leaves together with either GUS, AVR-Pii or AVR-PikD, and performed confocal microscopy as described in Methods. Interestingly, GFP:OsHIPP20 localized to dot-like structures in the cell wall. To further investigate these membrane puncta, we tested for co-localization with aniline blue, a known marker for callose deposition commonly associated with plasmodesmata (**Fig. 3B**; ^29^Thomas et al. 2008). Interestingly, when we co-expressed AVR-PikD with the HIPPs, GFP:OsHIPP19 and GFP:OsHIPP20 were relocalized from plasmodesmata to the cytosol (**Fig. 3B; Fig. S11**). Also, in the presence of GUS or AVR-Pii we found that GFP:OsHIPP19 and GFP:OsHIPP20 showed nucleo-cytoplasmic localization and accumulated in punctate structures with varying sizes (**Fig. 3C; Fig. S12**). When OsHIPP19 or OsHIPP20 was co-expressed with AVR-PikD, these punctae-like structures were not observed and the sHMA proteins were diffused in the cytoplasm and nucleus. This finding indicates that AVR-PikD binding can alter the subcellular distribution of sHMA proteins.

### *OsHIPP20* is a susceptibility gene to *Magnaporthe oryzae*

Of the seven sHMA proteins that interacted with AVR-PikD by Y2H, we selected OsHIPP19, OsHIPP20 and OsHPP04 for further study as were most frequently identified in this screen (**Table S1**), and showed strong interaction profiles with AVR-PikD (**Fig. 1**). To explore the function of these sHMA proteins in rice, we generated knockout (KO) mutants by CRISPR/Cas9-mediated mutagenesis in the rice cultivar Sasanishiki, which is susceptible to the blast fungus isolate Sasa2. We targeted *OsHIPP19*, *OsHIPP20* and *OsHPP04* (**Fig. 4A, Fig. S13**). The resulting KO lines were challenged with the *M. oryzae* isolate Sasa2. The KO lines of *OsHIPP19* (two independent lines) and *OsHPP04* showed a similar level of infection as the wild-type control (Sasanishiki) (**Fig. S13**). However, the *OsHIPP20* KO line showed a reduction in lesion size caused by *M. oryzae* infection (**Fig. 4**). In addition, a Sasanishiki line heterozygous for *OsHIPP20* wild-type and KO alleles were self-fertilized and its progeny were challenged with *M. oryzae* Sasa2. Progeny with homozygous *OsHIPP20* KO allele showed enhanced resistance as compared to their sib lines with *OsHIPP20* wild-type alleles, confirming that *OsHIPP20* is a susceptibility gene. We confirmed there was no off-target editing in other sHMA genes close to *OsHIPP20* by whole genome resequencing (**Table S2**). These results indicate that *OsHIPP20* is a susceptibility (*S-*) gene that is required for full infection of rice (cultivar Sasanishiki) by *M. oryzae*. Growth of *OsHIPP20*-KO Sasanishiki lines was comparable to the wild-type Sasanishiki (**Fig. 4B**; **Fig. S14**).

**Fig. 4.**
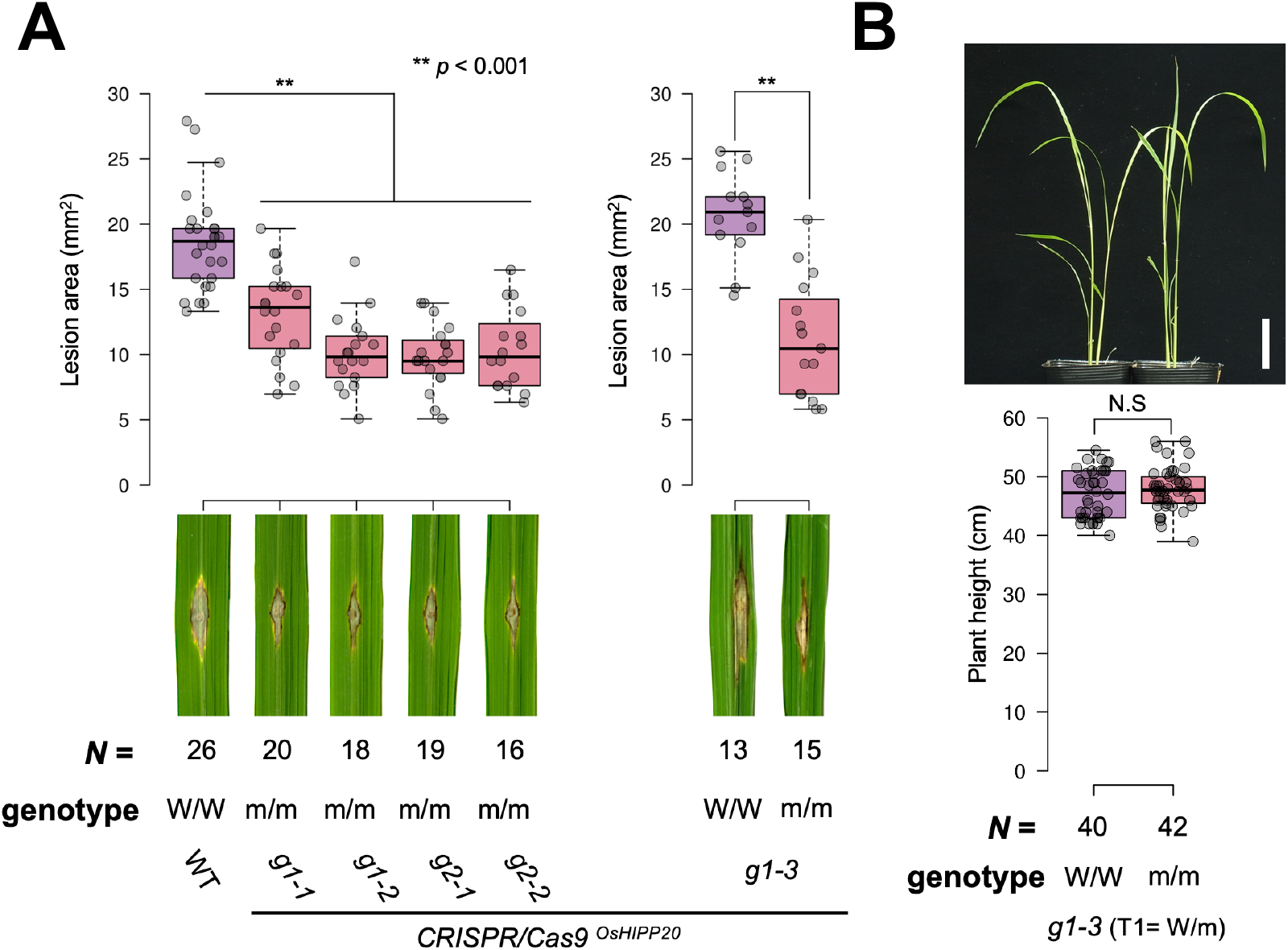
*OsHIPP20* is a susceptibility gene (S-gene). **A:** A compatible *Magnaporthe oryzae* isolate Sasa2 was punch inoculated onto the leaves of rice cultivar Sasanishiki as well as the *OsHIPP20*-knockout lines. Left: Punch inoculation results of T2 generation of two homozygous KO rice groups (guide RNA-1 [g1] and guide RNA-2 [g2], each with two replicates [g1-1, g1-2 and g2-1 and g2-2]). Right: Punch inoculation results of T2 progeny segregated to the wild type allele (W/W) and KO-type allele (m/m) from a T1 heterozygous KO line (g1-3). Box plots show lesion area sizes in the rice lines (top). Statistical significance is shown after t-test. Photos of typical lesions developed on the leaves after inoculation of *M. oryzae* (bottom). The number of leaves used for experiments are indicated below. **B:** No growth defect in *OsHIPP20* KO line as compared to the wild type one month after seed sowing. Top: Overview of the W/W and m/m plants. Bottom: Box plot showing plant height distribution of W/W and m/m plants segregated from the g1-3 heterozygous KO line. Scale bar: 5cm.

## Discussion

In this paper, we set out to identify host targets of the *M. oryzae* effector AVR-Pik, to explore the potential virulence function of this effector. We found that AVR-Pik binds multiple sHMA proteins of rice that belong to the same phylogenetic clade (Clade A), which also contain the integrated HMA domains of Pik-1 and RGA5 (**Fig. 1**). These findings support the view that NLR integrated domains have evolved from the host targets of pathogen effectors and that the HMA-containing proteins induced by chitin, a pathogen Pathogen-Associated Molecular Pattern (PAMP), are a major host target of plant pathogen effectors. In an independent study, ^27^Maidment et al. (2021) showed that AVR-Pik binds to OsHIPP19 with nanomolar affinity *in vitro* and showed the interaction of the effector with this sHMA is via an interface conserved with the Pik-1 integrated HMA domains providing further evidence that this effector targets host sHMA proteins.

Heavy metal-associated (HMA) domains were first defined in metal binding domains of P-type ATPase family copper transport proteins, including human MNK and WND proteins, mutations of which cause Menkes disease and Wilson disease, respectively (^30^Bull & Cox. 1994). HMA domains are also found in a number of heavy metal transport or detoxification proteins both in bacteria and eukaryotes. The yeast metallochaperone Atx1 was shown to deliver monovalent copper ions to the P-type ATPase Ccc2 that transports copper to trans-Golgi vesicle where it is taken up by the multicopper oxidase Fet3 (^31^Askwith et al. 1994; ^32^Pufahl et al. 1997; ^33^Rosenzweig & O’Halloran. 2000). A typical HMA domain contains two conserved cysteine residues involved in metal binding in a MxCxxC motif that is located towards the N-termini of the domain (^30^Bull & Cox, 1994).

In most organisms, only a small number of HMA-containing proteins have been reported. By contrast, in plants, proteins containing HMA-like domains have massively expanded (^34^Dykema et al. 1999; ^35^Barth et al. 2009; ^21^De Abreu-Neto et al. 2013). For example, ^35^Barth et al. (2009) identified 44 Arabidopsis genes that encode for proteins containing an HMA domain and a C-terminal putative isoprenylation motif (CaaX). Based on the presence or absence of the C-terminal isoprenylation motif, ^21^De Abreu-Neto et al. (2013) grouped plant sHMAs into heavy metal-associated isoprenylated plant proteins (HIPPs) and heavy metal-associated plant proteins (HPPs). We have chosen to use the naming convention of ^21^De Abreu-Neto et al. (2013) here. In this manuscript, we present an analysis of the HMA-like repertoire of the rice (cultivar Nipponbare) genome, revealing the presence of at least 87 HMA-containing small protein (abbreviated as sHMA) genes (**Fig. 1**).

The biological functions of plant sHMA proteins reported so far are diverse. Two Arabidopsis HMA-containing proteins, CCH and ATX1, complemented yeast *atx1* mutant, and are presumed to be involved in copper transport (^36^Himelblau et al. 1998; ^37^Puig et al. 2007; ^38^Shin et al. 2012). ^35^Barth et al. (2009) showed that the Arabidopsis HMA-containing protein HIPP26 localizes to nuclei and interacts with a zinc-finger transcription factor ATHB29, while ^39^Gao et al. (2009) reported the same protein (with an alternative name, ATFP6) was localized to plasma membrane and interacted with acyl-CoA–binding protein ACBP2, which was hypothesized to be involved in membrane repair after oxidative stress. ^40^Zhu et al. (2016) reported that the Arabidopsis HMA-containing protein NaKR1 interacts with Flowering Locus T (FT) and mediates its translocation from leaves to shoot apices. ^41^Cowan et al. (2018) reported that potato mop-top virus (PMTV) movement protein TGB1 interacts with *Nicotiana benthamiana* sHMA protein HIPP26 and relocalizes this protein from the plasma membrane to the nucleus, thus contributing to PMTV long-distance movement by altering transcriptional regulation.

Genetic studies have also revealed roles of specific plant sHMA proteins in defense and susceptibility towards pathogens. Deletion in the proline-rich domain of Pi21, a rice sHMA, conferred a partial resistance against compatible isolates of *M. oryzae* (^22^Fukuoka et al. 2009). Virus-induced gene silencing of wheat *TaHIPP1* enhanced resistance against stripe rust caused by *Puccinia striiformis* f. sp. *Triticii* (^42^Zhang et al. 2015). Similarly, a knockout mutant of *Arabidopsis AtHMAD1* enhanced resistance against virulent *Pseudomonas syringae* DC3000 (^43^Imran et al. 2016) and a knockout mutant of *Arabidopsis AtHIPP27* enhanced resistance against beet cyst nematode (^44^Rodakovic et al. 2018). However, it remains unclear how these sHMA proteins impact interactions with these diverse pathogens. Nonetheless, given that HMA domains have integrated into NLR immune receptors in at least four botanical families, it is likely that HMA containing proteins have repeatedly been targeted by pathogens across a diversity of flowering plant species and are thus important components in plant-pathogen interactions.

In addition to Pik-1, the NLR RGA5 also carries an integrated HMA domain that binds two *M. oryzae* effectors, AVR-Pia and AVR1-CO39. However, in our Y2H assays (with high stringency conditions), we didn’t detect any interaction between AVR-Pia and AVR1-CO39 and the tested sHMA proteins. We hypothesize that these two effectors may weakly bind the tested sHMAs or bind other rice sHMA proteins among the >80 members of this family.

In this study, we revealed that gene knockout of *OsHIPP20* confers enhanced resistance to rice against a compatible isolate of *M. oryzae* (**Fig. 4**). Therefore, like *Pi21*, *OsHIPP20* is a susceptibility gene (*S*-gene), whose activity is required for full virulence of the *M. oryzae* pathogen in rice. Rice *pi21* is established as a useful blast resistance gene (^22^Fukuoka et al. 2009). *OsHIPP20* knockout lines did not show growth defect. Therefore, combined with *pi21*, *oshipp20* mutants may provide a novel source of durable resistance against blast disease.

We also found that AVR-PikD binds and stabilizes OsHIPP19 and OsHIPP20 (**Fig. 3**). We hypothesize that AVR-Pik–mediated stabilization of sHMA proteins suppresses host defenses, resulting in enhanced *M. oryzae* invasion of rice cells (**Fig. 5**). Next, it will be important to determine the roles of the extended family of sHMA proteins in rice and other plants to understand the interplay between effector-mediated protein stabilization and disease.

**Fig. 5.**
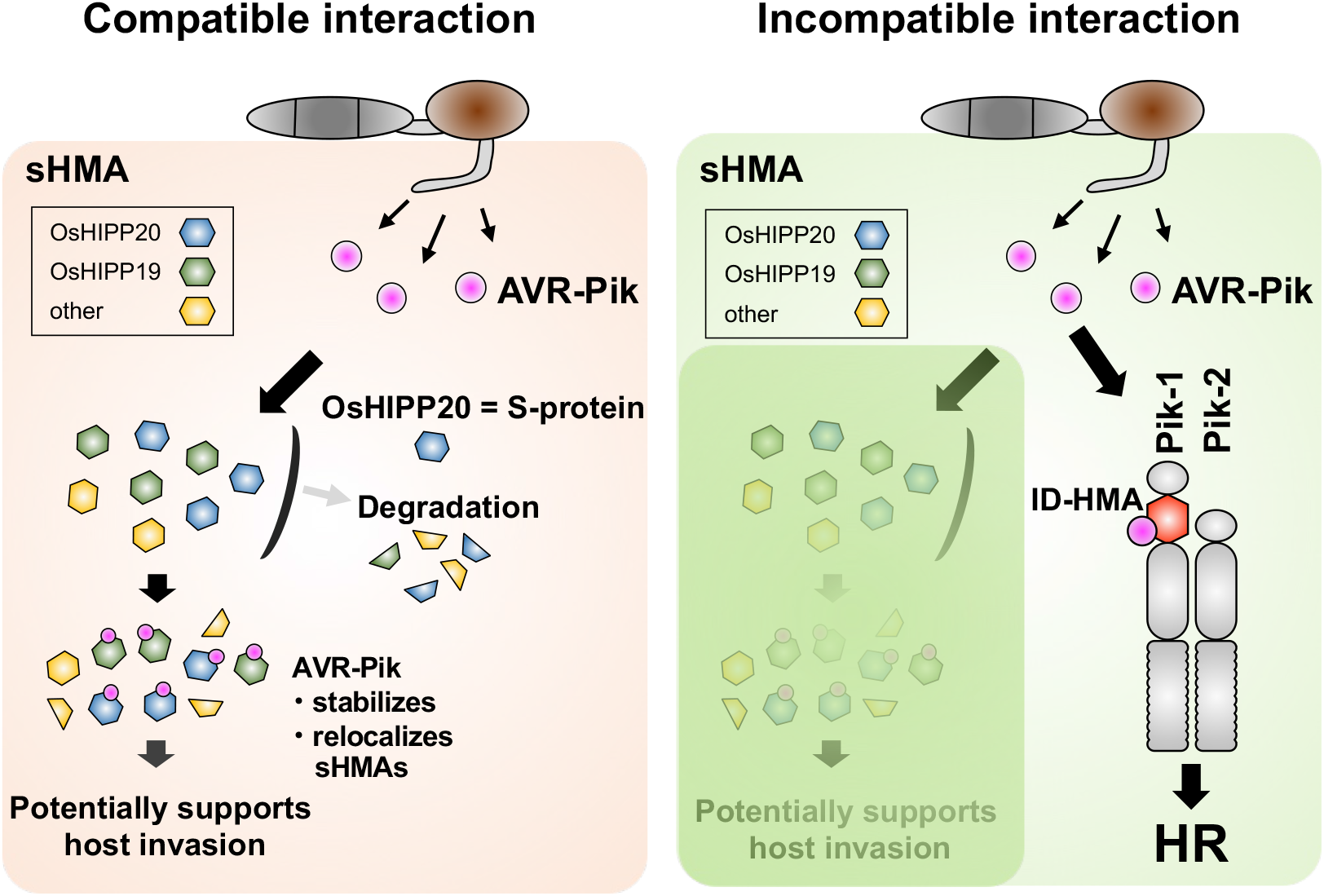
Schematic representation of a model showing molecular interactions between the AVR-Pik effector, rice sHMA proteins and Pik NLRs. In the compatible interaction (susceptible, left), AVR-Pik binds rice Clade A sHMA proteins, stabilizes and relocalizes them, possibly enhancing pathogen virulence. OsHIPP20 is a S-protein required for effective *M. oryzae* invasion. In the incompatible interaction (resistant, right), AVR-Pik interacts with integrated HMA domains of the Pik-1 NLRs which, together with Pik-2, triggers disease resistance by the hypersensitive response (HR). AVR-Pik and Pik seem involved in arms-race coevolution (selective force to enhancing interaction in Pik and evading interaction in AVR-Pik) by each generating multiple variants.

Our conceptual and mechanistic understanding of how plant NLR proteins perceive pathogens continues to expand. A model termed the integrated domain hypothesis postulates that NLRs can bait pathogen effectors directly through integrated decoy/sensor domains (^2^Cesari et al. 2014; ^3^Wu et al. 2015; ^4^Sarris et al. 2016; ^5^Kroj et al. 2016). These unconventional NLR domains are thought to have evolved by duplication and integration of an effector host target into the receptor protein. However, there are only very few examples where the evolutionary origin of the NLR integrated domain could be traced to an effector target (^45^Le Roux et al. 2015; ^46^Sarris et al. 2015; ^47^Grund et al. 2020). Here, we show that AVR-Pik interacts with sHMA proteins that belong to the same phylogenetic clade as the HMA domains integrated into the rice NLRs Pik-1 and RGA5. Therefore, throughout evolution, the Pik-1 NLR immune receptor has co-opted sHMA proteins through the integration of an HMA domain and neofunctionalization of this domain as a bait for the effector (**Fig. 5**). This has launched a coevolutionary arms race between Pik-1 and AVR-Pik. Given that binding of AVR-Pik to Pik-1 HMA domains is necessary for triggering cell death and disease resistance in rice, new variants of AVR-Pik have arisen in *M. oryzae* populations that evade binding the integrated Pik-1 HMA but maintain their virulence activity (^13^Kanzaki et al. 2012; ^12^Maqbool et al. 2015; ^16^Białas et al. 2018; ^48^Longya et al. 2019). Here we show that each of the AVR-Pik variants tested retain binding to OsHIPP19 and OsHIPP20 (**Fig. 1D**), consistent with the view that stealthy effectors can retain virulence activities. This demonstrates that effector variation can affect the phenotypic outcomes of disease susceptibility and resistance independently through mediating bespoke interactions with different HMA domains. This elegant model highlights a surprisingly intimate relationship between plant disease susceptibility and resistance, as well as microbial virulence, driven by complex coevolutionary dynamics between pathogen and host.

## Materials and Methods

### Construction of the maximum likelihood tree of HMA family genes

The protein sequences of the HMA domains were aligned by MAFFT (^49^Katoh et al. 2013) with the following method parameter set: --maxiterate 1000 --localpair. Then, the maximum likelihood tree was constructed by IQ-TREE (^50^Nguyen et al. 2015) with 1,000 bootstrap replicates (^51^Hoang et al. 2018). The model was automatically selected by ModelFinder (^52^Kalyaanamoorthy et al. 2017) in IQ-TREE (^50^Nguyen et al. 2015).

### RNA-seq of rice leaves and suspension cultured cells and gene expression analysis

Total RNA was extracted from rice leaves and cultured cells (one hour after mock or chitin treatment) using SV Total RNA Isolation System (Promega, United States). One microgram of total RNA was used to prepare sequencing libraries with NEB Next Ultra II Directional RNA Library Prep Kit for Illumina (NEB, United States). These libraries were sequenced by paired-end (PE) sequencing using the Illumina Hiseq platform (Illumina, United States). The quality of the RNA-seq was evaluated using the FastQC (http://www.bioinformatics.babraham.ac.uk/projects/fastqc/). After QC, filtered reads were used for further analysis. Hisat2 (^53^Kim et al., 2015) was used to align RNA-seq reads against the *O. sativa* reference genome downloaded from Rice Genome Annotation Project (http://rice.uga.edu/pub/data/Eukaryotic_Projects/o_sativa/annotation_dbs/pseudomolecules/). The levels of gene expression were scored by TPM (Transcripts Per Million) using StringTie (^54^Pertea et al. 2015).

### Yeast two-hybrid assay

To identify AVR-PikD–interacting proteins, signal peptide–truncated cDNA fragments of AVR-PikD were inserted into *Eco*RI and *Bam*HI sites of pGBKT7 (bait) vector (Takara Bio, Japan) to construct AVR-PikD/pGBKT7 (**Datasets S1**, ^13^Kanzaki et al. 2012). MATCHMAKER Library Construction & Screening kit (Takara Bio) was used to construct the rice cDNA library from leaf tissues of rice cultivar Sasanishiki 4, 24 and 48 h after inoculation with *M. oryzae* strain Sasa2 (race 037.1). Yeast strain AH109 competent cells were transformed with pGBKT7/AVR-PikD pGADT7-Rec and the rice cDNA library by using the polyethylene glycol/lithium acetate (PEG/LiAc) method, and plated on selective agar plates containing minimal medium without Trp, Leu, Ade and His, and supplemented with 20 mg/L of 5-Bromo-4-Chloro-3-indolyl a-D-galactopyranoside (X-α-Gal) and 10 mM 3-amino-1,2,4-triazole (3-AT). cDNAs in the library were transferred to pGAD-Rec vector harboring GAL4 activation domain (AD) by homologous recombination in yeast cells. Positive yeast transformants were streaked onto a minimal medium agar plate without Trp and Leu and used for sequence analysis. To examine the protein–protein interactions between sHMAs and AVR-Pia, AVR-Pii, AVR1-CO39 and AVR-Pik alleles, yeast two-hybrid assay was performed as described previously (^13^Kanzaki et al. 2012). Bait and prey plasmid vectors were constructed as described in **Datasets S1**. Signal peptide–truncated cDNA fragments of AVRs were amplified by PCR by using primer set (**Datasets S1**) and inserted into *Eco*RI and *Bam*HI sites of pGADT7 (prey) or pGBKT7 (bait) vectors (Takara Bio). sHMA cDNAs were synthesized from total RNAs of rice leaves (cultivar Sasanishiki) and inserted into pGADT7 and pGBKT7 by using *Spe*I and *Bam*HI sites as described in **Datasets S1**. In the case of sHMAs containing *Spe*I or *Bam*HI site, In-Fusion HD Cloning Kit (Takara Bio) was utilized to construct plasmid vectors. The various combinations of bait and prey vectors were transformed into yeast strain AH109 by using the PEG/LiAc method. To detect the protein–protein interactions, ten-fold dilution series (×1, ×10^-1^, ×10^-2^) of yeast cells (×1 : OD600=1.0) were spotted onto on basal medium lacking Trp, Leu, Ade and His but containing X-α-Gal (Takara Bio) and 10 mM 3-amino-1,2,4-triazole (3-AT). Positive signals were evaluated by blue coloration and growth of the diluted yeast. As a control, yeast growth on basal medium lacking Trp, Leu was also checked. Details of plasmids used are indicated in **Datasets S1**. To check the protein accumulation in yeast cells, each transformant was propagated in the liquid basal medium lacking Trp, Leu with gentle shaking at 30℃ overnight. Yeast cells from 10 ml medium were collected and 100 mg of yeast cells were treated with 400µl of 0.3 N NaOH for 15 min at room temperature. Resulting yeast extracts were used for western blot analysis using anti-Myc HRP-DirecT (MBL, Japan) for bait proteins and anti-HA (3F10)-HRP (Roche, Switzerland) for prey proteins.

### Plasmid construction for AlphaScreen

A total of 18 sHMAs belonging to CladeA and CladeB (Fig. 1AB) were amplified by PCR by using primer sets (**Datasets S1**) and inserted into *Bsa*I sites of the level 0 vector pICH41308 (Addgene no. 47998) for the Golden Gate cloning (^55^Engler et al. 2008). FLAG-tagged sHMA was generated by Golden Gate assembly with pICH45089 (35S promoter (double), Addgene no. 50254), pAGT707 (5’ U-TMV+Ω, Addgene no. 51835), pICSL30005 (3xFLAG, Addgene no. 50299), pICH<u>41308::sHMA</u>and pICH41414 (3’UTR+ terminator, Addgene no. 50337) into a binary vector pICH47732 (Addgene no. 48000). Using pICH47732::sHMA as the PCR template, FLAG-tagged sHMA was amplified with forward primer (5’-CTACATCACCAAGATATCATGGATTATAAGGACCATGA-3’) and reverse primer (5’-TCTATACAAAACTAGTACTCACACATTATTATGGAG-3’), and then cloned into pEU-E01 vector (^56^Sawasaki et al. 2002) for wheat cell-free protein synthesis.

### Preparation of linier DNA templates for in vitro transcription and wheat cell-free protein synthesis

The *in vitro* transcription templates for the FLAG tagged sHMA were prepared by standard PCR with primer pairs (Spu, 5’- GCGTAGCATTTAGGTGACACT- 3’; SP-A1868, 5’-CCTGCGCTGGGAAGATAAAC -3’) using pEU-E01::sHMA expression plasmids as templates (as mentioned above). For preparation of *in vitro* transcription template for biotin-Myc tagged AVR-PikD was amplified by a slightly modified two-step PCR method (^56^Sawasaki et al. 2002) using pGBKT7::AVR-PikD plasmid as template. In the first round of PCR, the gene containing ORF and Myc-tagged sequence region was amplified by PCR using following primer pairs: AVR-PikD-S1, 5’- ccacccaccaccaccaATGGAGGAGCAGAAGCTGATCTC -3’(lower case letters indicate S1-linker sequence); AODA2306, 5’- AGCGTCAGACCCCGTAGAAA -3’. Next, a second round of PCR was carried out to add the SP6 promoter, the translation enhancer sequence E01, and the biotin ligation site sequence at the 5’ end of the ORF using the first PCR product as template. The following two sense primers and one antisense primer were used for PCR: Spu, 5’- GCGTAGCATTTAGGTGACACT -3’; deSP6E02-bls-S1, 5’- GGTGACACTATAGAACTCACCTATCTCTCTACACAAAACATTTCCCTACATACAACTTTC AACTTCCTATTATG<u>GGCCTGAACGACATCTTCGAGGCCCAGAAGATCGAGTGGCACGAA</u> CTccacccaccaccaccaATG -3’ (under line and lower case letters indicate the biotin ligation site and S1-linker sequence, respectively); AODA2303, 5’- GTCAGACCCCGTAGAAAAGA -3’. All amplified PCR products were confirmed by agarose gel electrophoresis, and used for *in vitro* transcription and wheat cell-free protein synthesis.

### Cell-free protein synthesis

In vitro transcription and wheat cell-free protein synthesis were performed using WEPRO7240 expression kit (Cell-Free Sciences, Japan). Transcript was made from each of the DNA templates mentioned above using the SP6 RNA polymerase. The translation reaction was performed in the bilayer mode (^57^Takai et al. 2010) using WEPRO7240 expression kit (Cell-Free Sciences) according to the manufacture’s instruction. For biotin labeling, 1 μl of crude biotin ligase (BirA) produced by the wheat cell-free expression system was added to the bottom layer, and 0.5 μM (final concentration) of d-biotin (Nacalai Tesque, Japan) was added to both upper and bottom layers, as described previously (^58^Sawasaki et al. 2008). The aliquots were used for the expression analysis and protein-protein interaction assay.

### Protein-protein interaction assays using AlphaScreen

The protein-protein interaction (PPI) between biotinylated AVR-PikD and FLAG-tagged sHMA proteins were detected with AlphaScreen technology provided by PerkinElmer. PPI assays were carried out in a total volume of 15 μl containing 1 μl of biotinylated AVR-PikD, and 1 μl of FLAG- sHMA proteins in the AlphaScreen buffer (100 mM Tris-HCl (pH8.0), 0.01% Tween20, 1mg/ml BSA) at 25 °C for 1 h in a 384-well Optiplate (PerkinElmer, San José, USA). In accordance with the AlphaScreen IgG (ProteinA) detection kit (PerkinElmer) instruction manual, 10 μl of detection mixture containing AlphaScreen buffer, 5 μg/mL anti-DYKDDDDK monoclonal antibody (clone 1E6, FUJIFILM Wako, Japan), 0.1 μl of streptavidin-coated donor beads, and 0.1 μl of Protein A-coated acceptor beads were added to each well of the 384-well Optiplate, followed by incubation at 25 °C for 1 h. Luminescence was analyzed using the AlphaScreen detection program using EnSight Multimode Plate Reader (PerkinElmer). All data represent the average of three independent experiments and the background was controlled using a dihydrofolate reductase (DHFR) from *E. coli*.

### Binding complex modeling by ColabFold

For the prediction of AVR-PikD/sHMA complex structures, we used the alphafold2advanced.ipynb notebook (ColabFold_v1.5.2) (^28^Mirdita et al., 2021) (https://colab.research.google.com/github/sokrypton/ColabFold/blob/v1.5.2/AlphaFold2.ipynb, accessed on August to October 2023) with the default mode. Display of the predicted AVR-PikD/sHMA. complex and measurement of interatomic distances were performed using Waals (Altif Laboratories Inc., Tokyo, Japan). Multimer confidence score is calculated as AlphaFold-multimer score (0.8*ipTM+0.2*pTM) in **Fig. S5, S6** (^59^Yin et al. 2022).

### Generation of rice mutants of *OsHIPP*s by CRISPR/Cas9 system

Rice plants with mutated *OsHIPP19*, *OsHIPP20* or *OsHPP04* were generated using the CRISPR/Cas9 system developed by ^60^Mikami et al. (2015). Primers OsHIPP19_gRNA1-F: 5’- gttgAAGCTGGTGGTGATGGCCTC-3’ and OsHIPP19_gRNA1-R: 5’-aaacGAGGCCATCACCACCAGCTT-3’ were annealed and cloned into pU6::ccdB::gRNA cloning vector by digestion with BbsI as the target sequence. The target sequence with the OsU6 promoter was replaced into the pZH::gYSA::MMCas9 vector by digestion with *Asc*I and *Pac*I, generating pZH::gYSA::MMCas9::OsHIPP19_gRNA1. Binary vector, pZH::gYSA::MMCas9:: OsHIPP19_gRNA2, pZH::gYSA::MMCas9::OsHIPP20_gRNA1, pZH::gYSA::MMCas9::OsHIPP20_gRNA2 and pZH::gYSA::MMCas9:: OsHPP04_gRNA1 were constructed by the same method; details of the primers are listed in **Datasets S1**. The rice cultivar ‘Sasanishiki’ was used for Agrobacterium-mediated transformation following the methods of ^61^Toki et al. (2006). Thereafter, regenerated T0 plants were sequenced using primers listed in **Datasets S1** and the mutation type was analyzed.

### Rice pathogenicity assays

Rice leaf blade punch inoculations were performed using the *M. oryzae* strains Sasa2 (without AVR-Pik alleles). A conidial suspension (3 × 10^5^ conidia mL^-1^) was punch inoculated onto the rice leaf one month after sowing. The inoculated plants were placed in a dew chamber at 27°C for 24 h in the dark and transferred to a growth chamber with a photoperiod of 16 h. Disease lesions were scanned 10 days post-inoculation(dpi) and the lesion size was measured using ‘Image J’ software (^62^Schneider et al. 2012).

### Transient gene expression assay in *N. benthamiana*

For transient protein expression, *Agrobacterium tumefaciens* strain GV3101 was transformed with the relevant binary constructs (**Datasets S1**). To detect sHMA (OsHIPP19 and OsHIPP20) protein accumulation and stability in *N. benthamiana*, several combinations of *Agrobacterium* transformants (the ratio of each transformant is 2:7:1 for sHMA:AVR (GUS) : P19; final concentration is OD_600_=1.0) were infiltrated using a needleless syringe into leaves of 3- to 4-weeks-old *N. benthamiana* plants grown at 23℃ in a greenhouse. Two days after infiltration, leaves were collected and homogenized by using a Multi-Beads Shocker (Yasui-Kikai, Japan) under cooling with liquid nitrogen. Then 2 ml of extraction buffer (10% glycerol, 25 mM Tris–HCl pH 7.0, 10 mM DTT, 1tablet / 50ml cOmplete™ Protease Inhibitor Cocktail [Roche, Switzerland]) was added to 1 mg of leaf tissues and further homogenized. After centrifugation at 20,000×g for 15 min, the supernatant was collected and the pellet was resuspended in 2 ml (the same as supernatant volume) of extraction buffer. The supernatants and pellet samples were subjected to SDS-PAGE followed by western blotting. Proteins were immunologically detected by using anti-HA (3F10)-HRP (Roche, Switzerland), anti-Myc-tag (HRP- DirecT) (MBL, Japan) and Anti-β-Glucuronidase (N-Terminal) antibodies produced in rabbit (Sigma-Aldrich, United States). The luminescent images were detected by luminescent Image Analyzer LAS-4000 (Cytiva, Japan) after treatment of ChemiLumi One Super or Ultra (Nacalai Tesque, Japan).

### Co-immunoprecipitaion assay

For co-immunoprecipitation assays, Myc-tagged sHMA (eGFP) and HA-tagged AVR-PikD binary constructs (**Datasets S1**) were transformed into *Agrobacterium tumefaciens* strain GV3101. Proteins were co-expressed in *N. benthamiana* and extracted from the leaves (approximately 150 mg) with 400 mL of extraction buffer (50 mM Tris-HCl pH 7.5 and 150 mM NaCl). Extracts were incubated with Anti-Myc-tag mAb-Magnetic Agarose (MBL, Japan) at 4°C for 1 h. Myc-agarose was washed with the same buffer 3 times and bound protein was eluted with 1x SDS sample buffer. The eluates were used for western blot analysis using anti-HA (3F10)-HRP (Roche, Switzerland) and anti-Myc-tag (HRP-DirecT) (MBL) antibodies.

### Localization of OsHIPP19 and OsHIPP20

To visualize subcellular localization of OsHIPP17, OsHIPP19 and OsHIPP20, GFP-tagged OsHIPP17 , OsHIPP19, OsHIPP20 expression constructs were generated by Golden Gate assembly with pICH45089, pAGT707, pICH41531 (GFP, Addgene no. 50321), pICH41308::OsHIPP17 (::OsHIPP19, ::OsHIPP20) and pICH41414 into a binary vector pICH47732. These constructs were transformed into *Agrobacterium tumefaciens* strain GV3101 for transient expression in *N. benthamiana* (the ratio of each transformant is 2:7:1 for GFP : AVR (GUS) : P19; final concentration is OD_600_=1.0). To visualize plasmodesmata, Aniline blue fluorochrome (Biosupplies, Australia) was used at 0.01 mg/ml and infiltrated into the leaves 2 days after agroinfiltration before being analyzed by confocal microscopy (^29^Thomas et al. 2008). The fluorescence was visualized by a Nikon AX Confocal Microscope System (Nikon, Japan). Aniline blue fluorochrome was excited using 405 nm laser and captured at 460-480 nm. GFP was excited using a 488 nm laser and captured at 505–555 nm (Fig. 3B).

For studying subcellular localization of OsHIPP19 and OsHIPP20 in Fig. 3C, pCambia::3xMyc-eGFP-OsHIPP19 and pCambia::3xMyc-eGFP-OsHIPP20 were transformed into *Agrobacterium tumefaciens* strain GV3101 for transient expression in *N. benthamiana* (the ratio of each transformant is 2:7:1 for GFP : AVR (GUS) : P19; final concentration is OD_600_=1.0). GFP fluorescence in the leaves 2 days after agroinfiltration were observed by an Olympus FluoView FV1000-D confocal laser scanning microscope (Olympus). GFP was excited with an HeNe(G) laser. A DM488/543/633 diachronic mirror, SDM beam splitter, and BA505-525 emission filter were used for observation.

## Acknowledgements

This work was supported by JSPS Grant 15H05779, 20H05681, 23K20042 to RT, 21K14834 to MS, and 18K05657 to HS; a Grant from JSPS/The Royal Society Bilateral Research for the project “Retooling rice immunity for resistance against rice blast disease” (2018-2019); the UKRI Biotechnology and Biological Sciences Research Council (BBSRC) Norwich Research Park Biosciences Doctoral Training Partnership, UK [grant BB/M011216/1]; the UKRI BBSRC, UK [grants BB/W00108X/1, BB/P012574, BB/M02198X]; the European Research Council [ERC; proposal 743165]; the Gatsby Charitable Trust; and the John Innes Foundation. We thank Dr. Yukio Kawamura, Iwate University for technical supports.

## Author Contributions

Author contributions: K.O., S.K. and R.T. designed research; K.O., K.F., M.S., T.T., K.N., H.S., A.H. N.M., A.B., T.L., R.K., S.K. and R.T. performed research; K.O., K. F., M.S., T.T., H.S., A.B., T.L., T.O.B., S.C., T.K., M.J.B., S.K. and R.T. analyzed data; and K.O., K.F., M.S., M.J.B., S.K., and R.T. wrote the paper.

## Supporting Information for

**Table S1.** Rice proteins that interacted with AVR-PikD in the Y2H screen. We carried out two Y2H screens (1^st^ and 2^nd^). The number of positive clones with insert sequences corresponding to the designated proteins are shown.

| Gene name (RGAP) | Gene Product | From De Abreu-Neto et al. | 1st screening |  | 2nd screening |  | total |  |
| --- | --- | --- | --- | --- | --- | --- | --- | --- |
| LOC_Os02g35329 | RING finger | - | 8 |  | 8 |  | 16 |  |
| LOC_Os04g39350 | Heavy metal associated (HMA) domain containing | OsHIPP19 | 1 |  | 4 |  | 5 |  |
| LOC_Os04g39010 |  | OsHIPP20 | 1 |  | 2 |  | 3 |  |
| LOC_Os02g37300 |  | OsHPP04 | 1 | **total** | 1 | **total** | 2 | **total** |
| LOC_Os02g37290 |  | OsHPP03 | 0 | 3 | 1 | 8 | 1 | 11 |
| LOC_Os01g22490 | Ubiquitin | - | 7 |  | 2 |  | 9 |  |
| LOC_Os09g25060 | WRKY76 | - | 5 |  | 1 |  | 6 |  |
| LOC_Os12g38170 | PR-5 | - | 0 |  | 5 |  | 5 |  |
| LOC_Os04g33390 | Unknown | - | 3 |  | 2 |  | 5 |  |

**Fig. S1.**
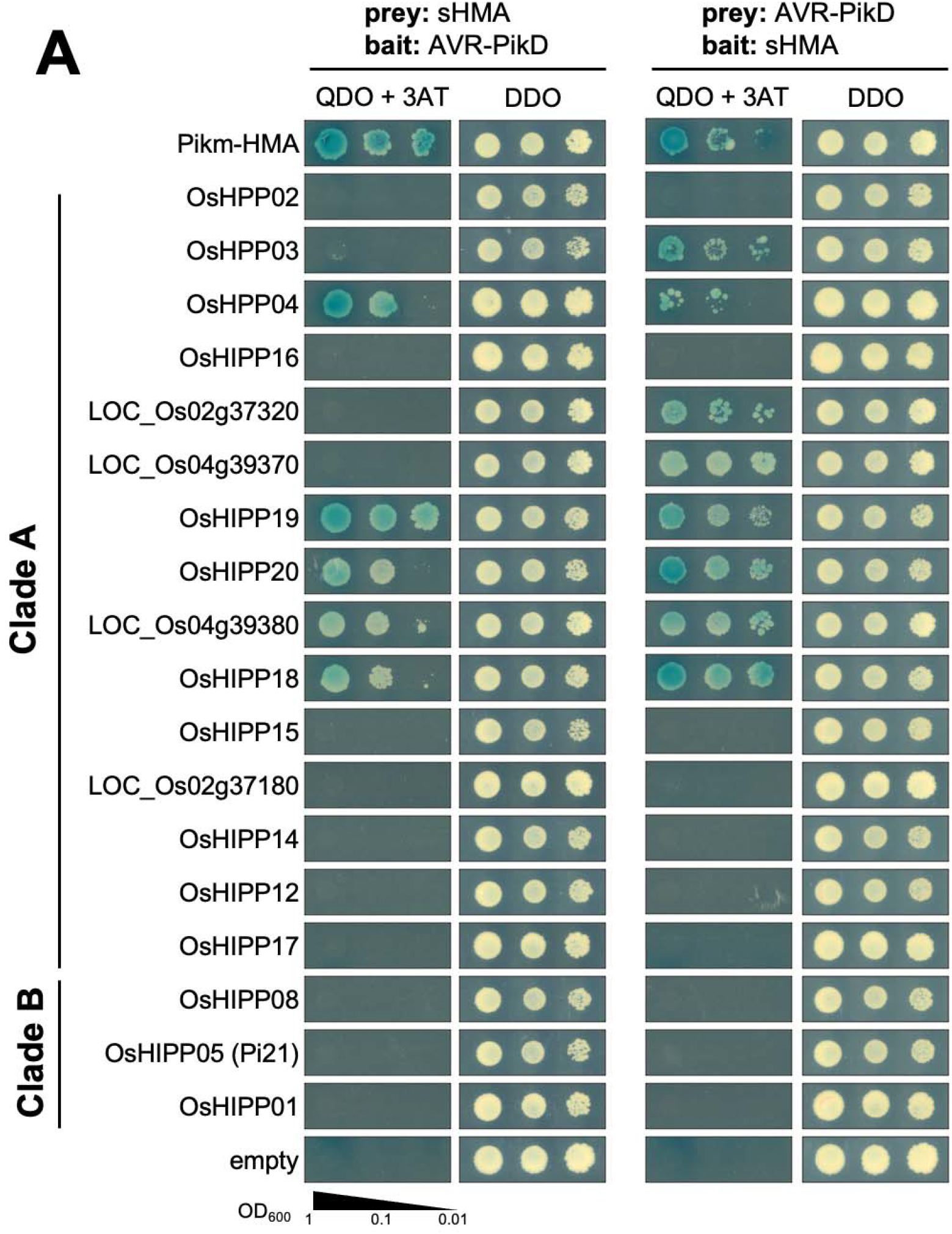
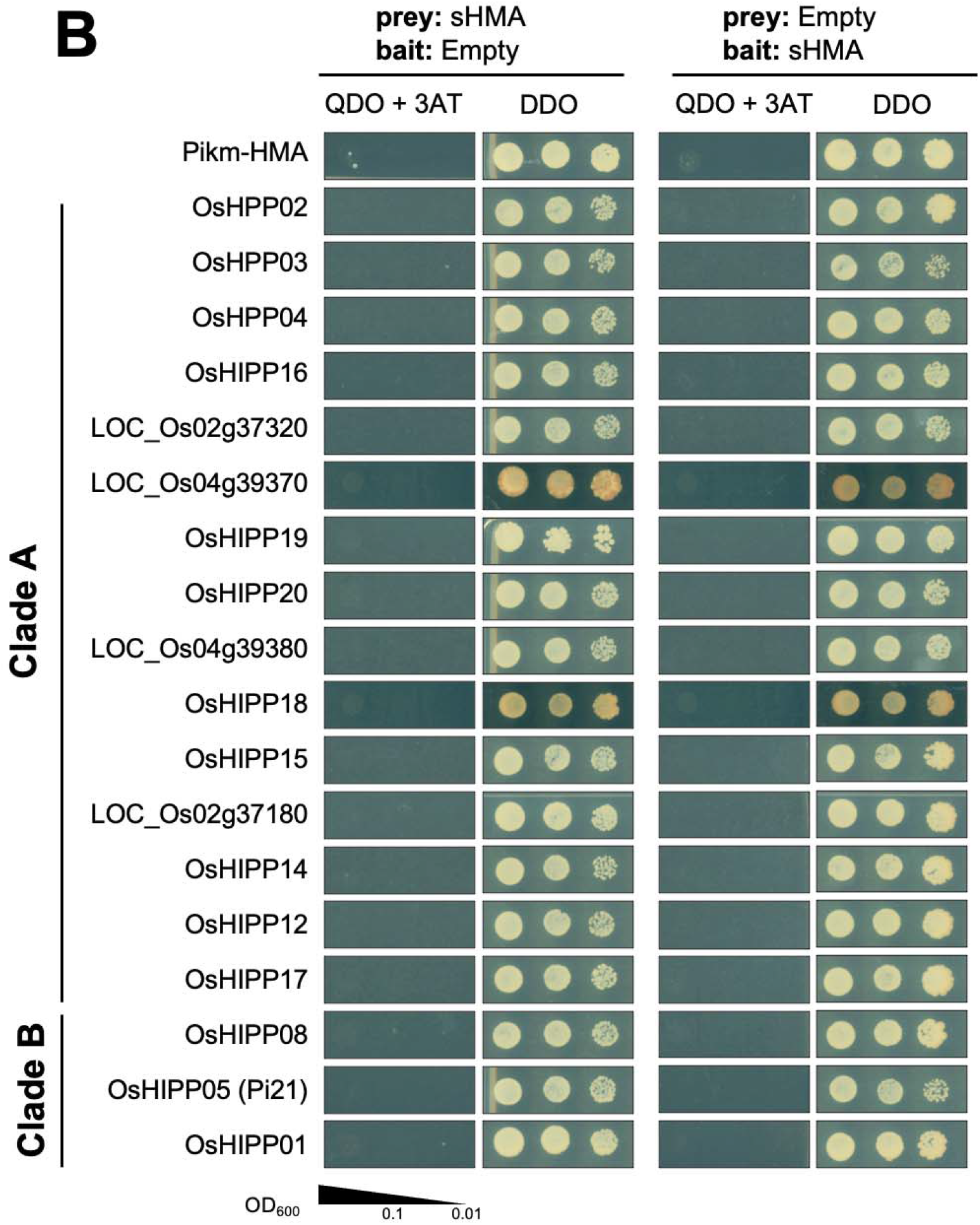
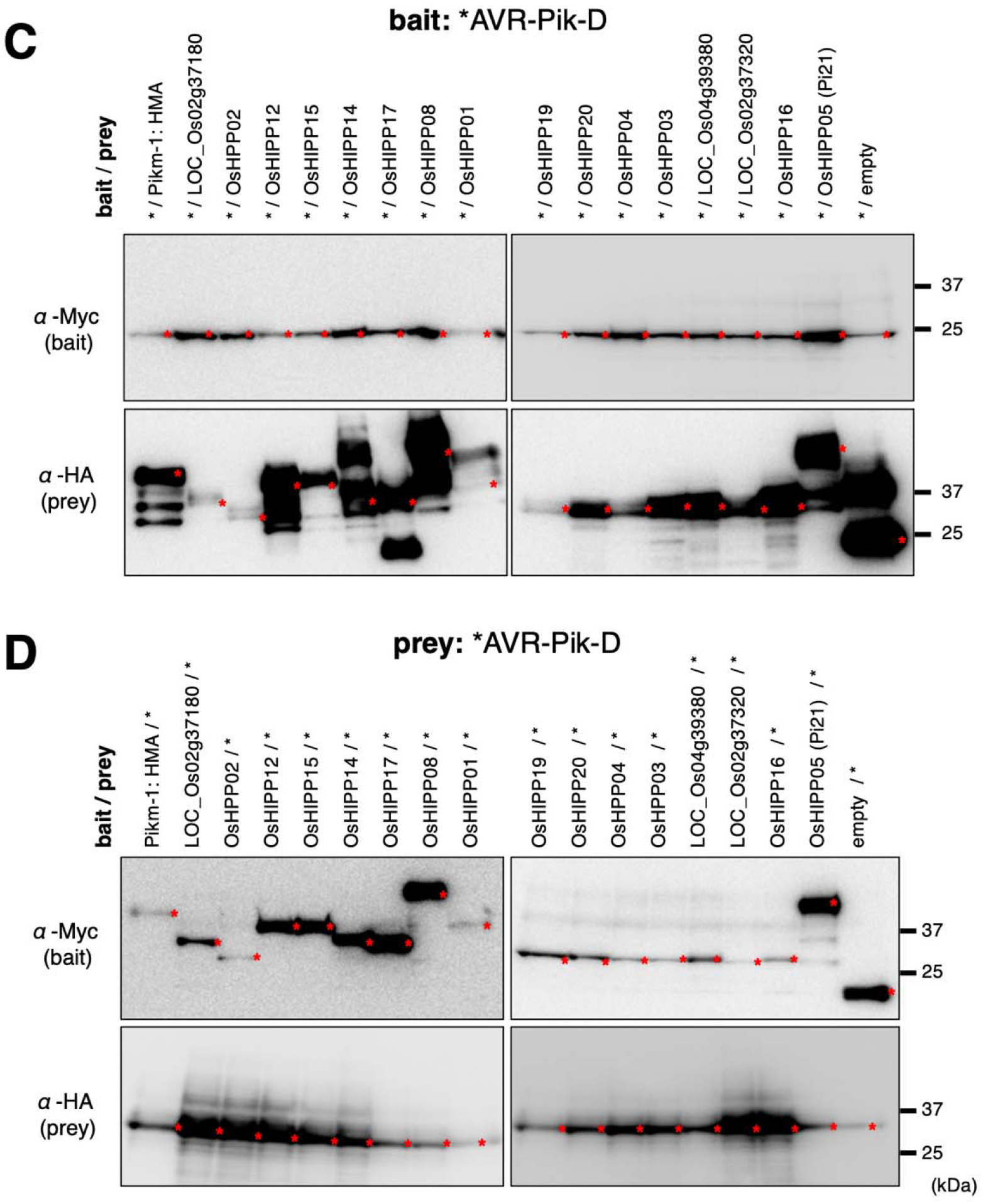
AVR-PikD interacts with Clade A sHMA proteins in yeast 2-hybrid assay (Y2H). **A:** Interactions between AVR-PikD and a subset of sHMA proteins were tested by Y2H. sHMA proteins were used as prey and AVR-PikD as bait (left panels) and AVR-PikD was used as prey and sHMA proteins as bait (right panels). Results with the conditions of stringent selection (QDO+3AT: SD/-Trp/-Leu/-Ade/- His, X-α-Gal,10mM 3AT) as well as no selection (DDO: SD/-Trp/-Leu) are shown. **B:** Interactions between empty vector products and a subset of sHMA proteins were tested by Y2H. sHMA proteins were used as prey and empty vector product as bait (left panels) and empty vector product was used as prey and sHMA proteins as bait (right panels). Results with the conditions of stringent selection (QDO+3AT) as well as no selection (DDO) are shown. **Western blot analysis confirms protein production in the Y2H experiment shown in A. C:** AVR-Pik-D was used as bait and sHMA proteins as prey. **D:** AVR-Pik-D was used as prey and sHMA proteins as bait. The bait protein was tagged with the Myc epitope and the prey protein tagged with the HA epitope. The protein bands expressed from each vectors are marked by red asterisks. The positions of molecular size marker are indicated in the right (kDa).

**Fig. S2.**
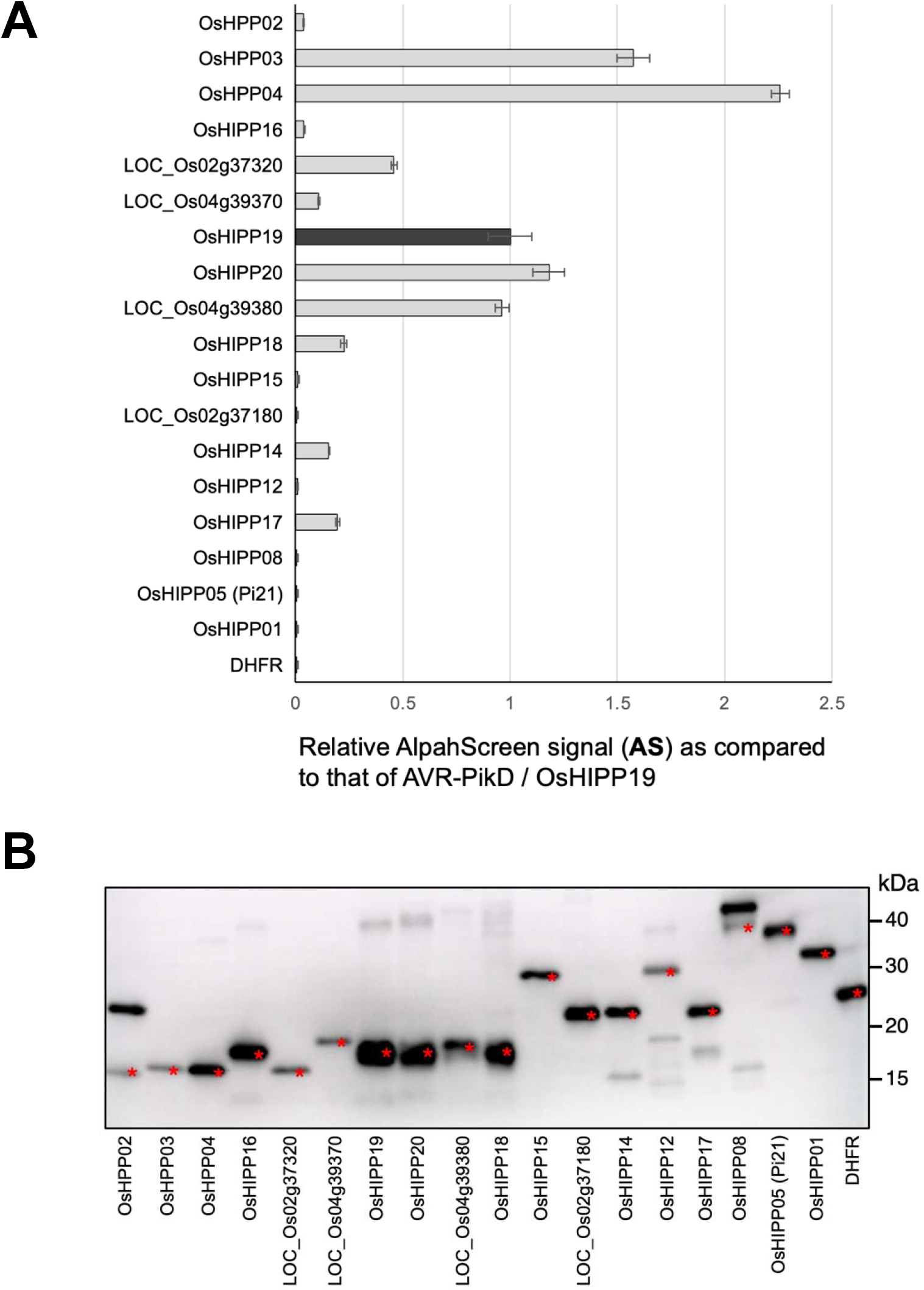
Interaction between AVR-PikD and Clade A sHMA proteins as addressed by AlphaScreen. **A:** AVR-PikD and sHMA proteins were produced by wheat germ translation system, and were subjected to AlphaScreen interaction assay. The values indicate relative AlpahScreen signal (**AS**) as compared to that of AVR-PikD / OsHIPP19. **B:** Western blot analysis confirms protein production in the AlphaScreen as shown in **A**. The sHMA proteins were tagged with the FLAG epitope and detected by anti-FLAG antibody. The protein bands expressed from the vectors are marked by red asterisks. The positions of molecular size marker are indicated on the right (kDa).

**Fig. S3.**
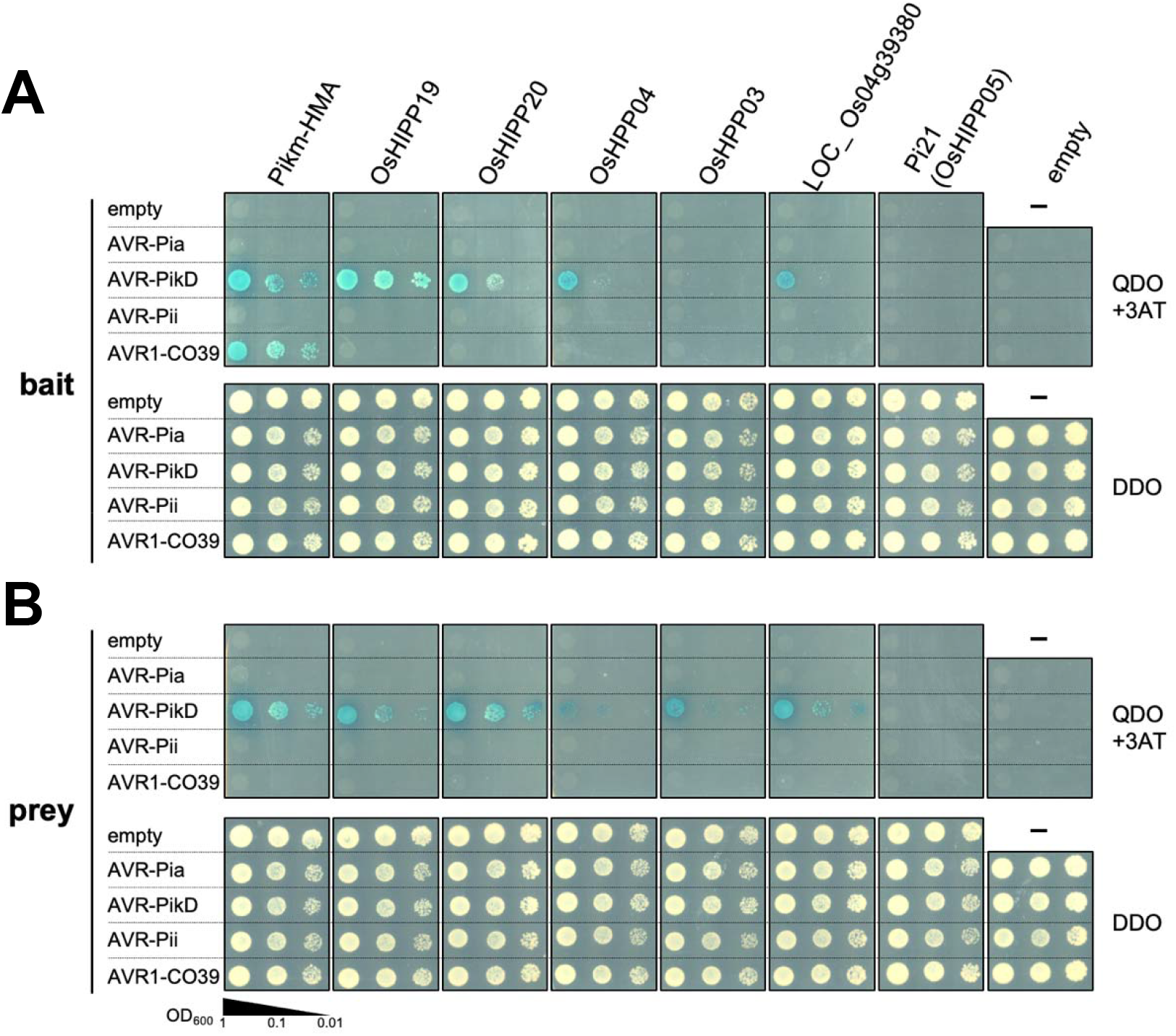
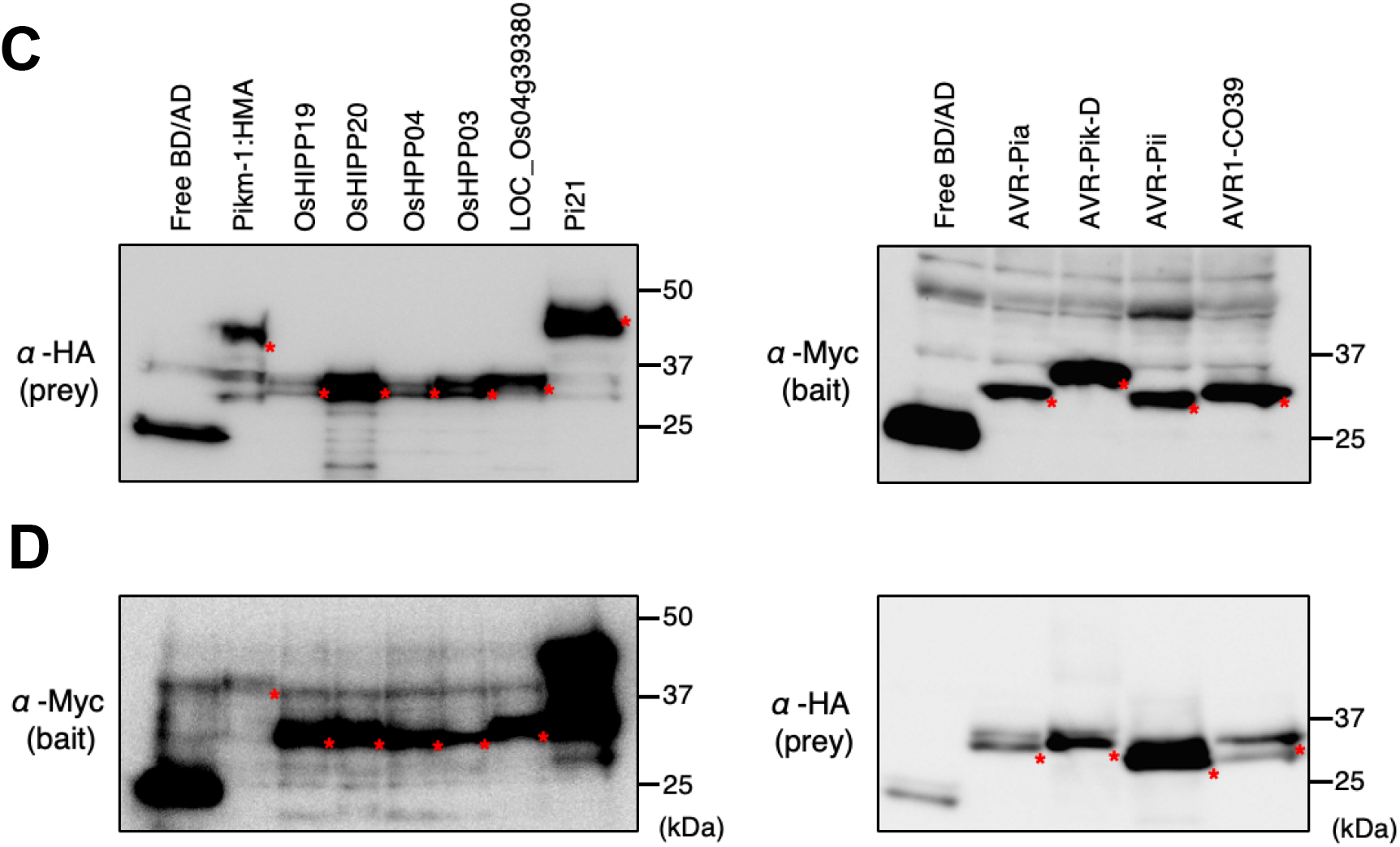
AVR-Pia and AVR1-CO39 do no bind Clade A sHMAs or Pi21. **A:** Four *Magnaporthe oryzae* effectors, AVR-Pia, AVR-PikD, AVR-Pii and AVR1-CO39 were tested for their binding with Clade A sHMAs (OsHIPP19, OsHIPP20, OsHPP04, OsHPP03 and LOC_Os04g39380) as well as Pi21 (OsHIPP05) of Clade B in Y2H assay with high stringency condition (QDO+3AT) as well as no selection (DDO). The HMA domain of Pikm-1 NLR protein (Pikm-HMA) interacts with AVR-PikD (Kanzaki et al. 2012) and used as a positive control. AVR-Pii was used as a negative control. Top panels **A** show the results when effectors were used as bait and sHMAs as prey, bottom panels **B** show the results when effectors were used as prey and sHMAs as bait. **Western blot analysis confirms protein production in Y2H experiment as shown in A and B. C:** Western blot results corresponding to Y2H in **A**. Pikm-1-HMA as well as sHMAs (prey) were detected by anti-HA antibody (left panel), whereas AVRs (bait) were detected by an anti-Myc antibody (right panel). **D:** Western blot results corresponding to Y2H in **B**. Pikm-1-HMA as well as sHMAs (bait) were detected by anti-Myc antibody (left), whereas AVRs (prey) were detected by an anti-HA antibody (right). The protein bands expressed from the constructs were marked by red asterisks. Molecular sizes (kDa) are indicated in the right of panels.

**Fig. S4.**
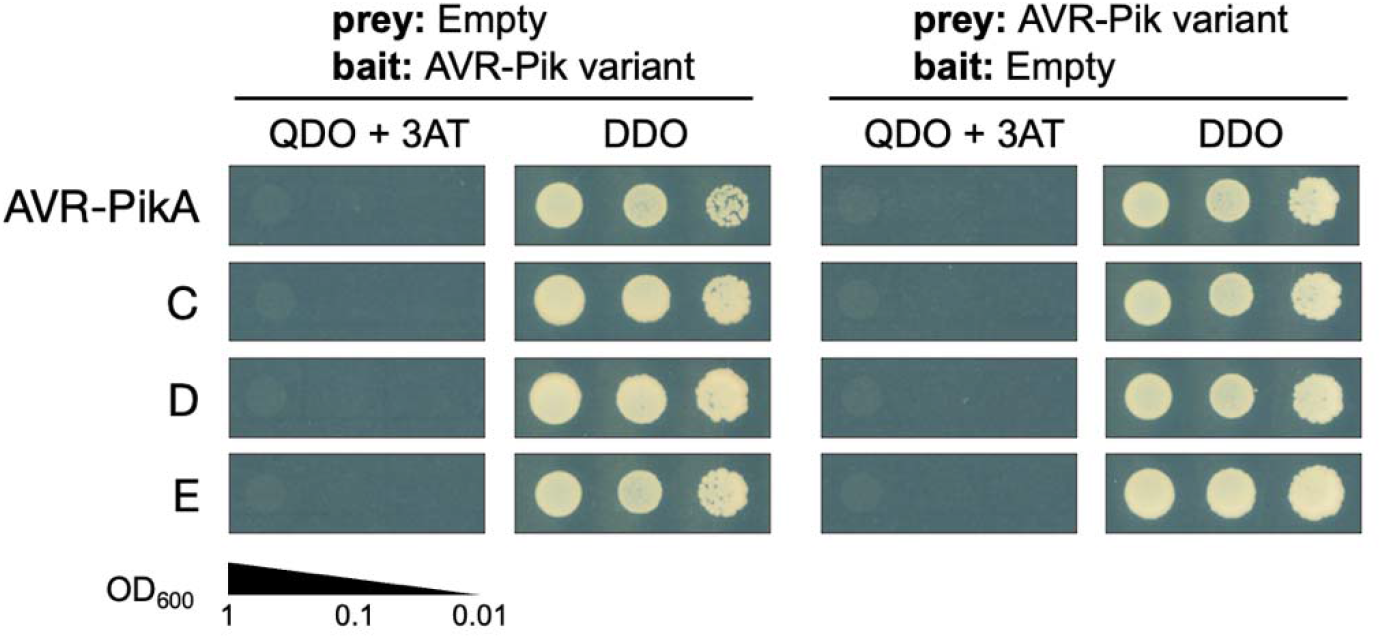
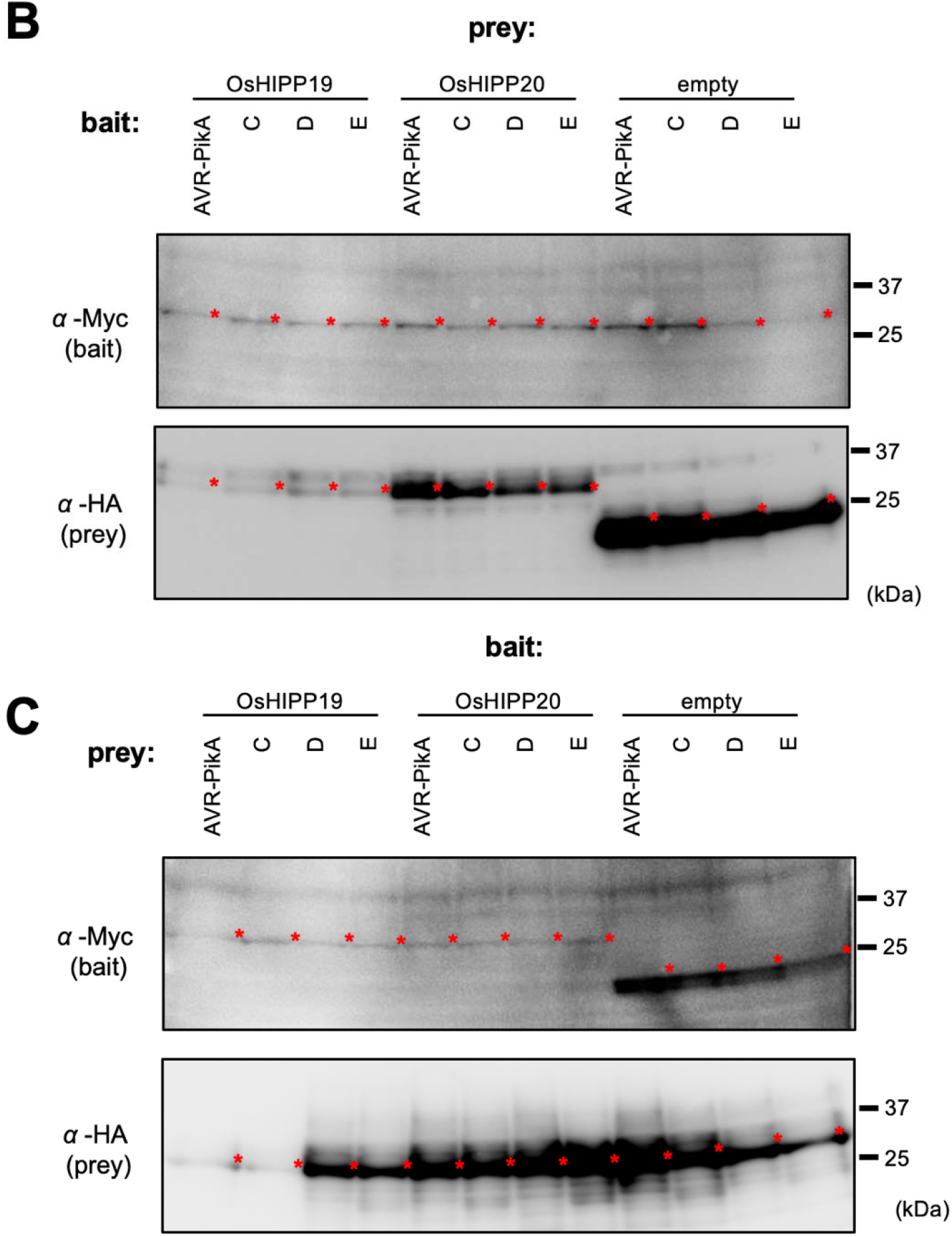
Y2H results with empty vector indicate no autoactivation with the AVR-Pik effector constructs. Results of high stringency selection (QDO+3AT) as well as no selection (DDO) are shown. **Western blot analysis confirming AVR-Pik-alleles (A, C, D, E) and OsHIPP19 and OsHIPP20 protein production in Y2H experiment as shown in Fig. 1D B:** OsHIPP proteins were used as prey and AVR-Pik alleles as bait. **C:** OsHIPP proteins were used as bait and AVR-Pik alleles as prey. Protein bands expressed from each vectors were marked by red asterisks. The positions of molecular size marker were indicated at right (kDa).

**Fig. S5.**
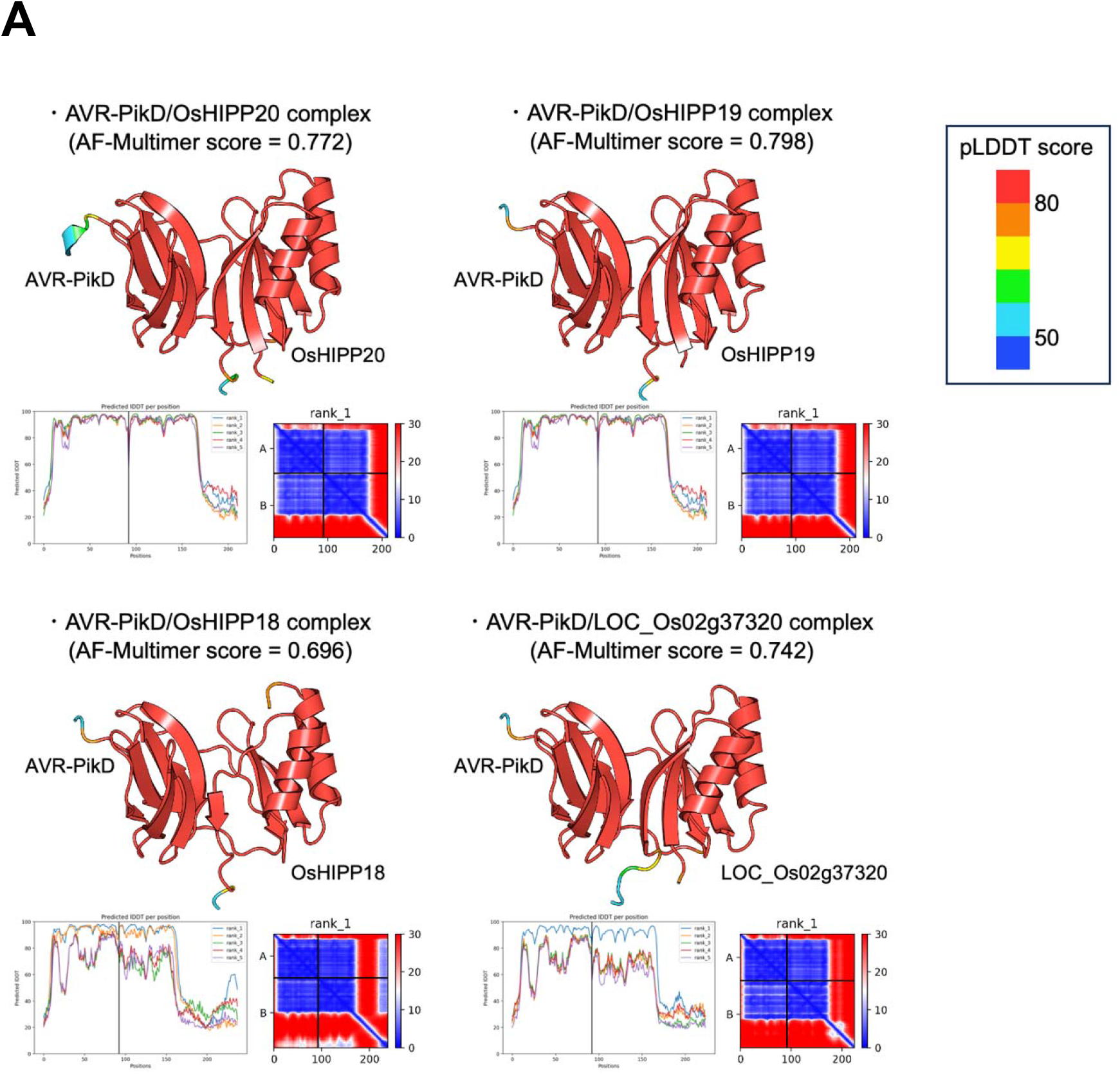
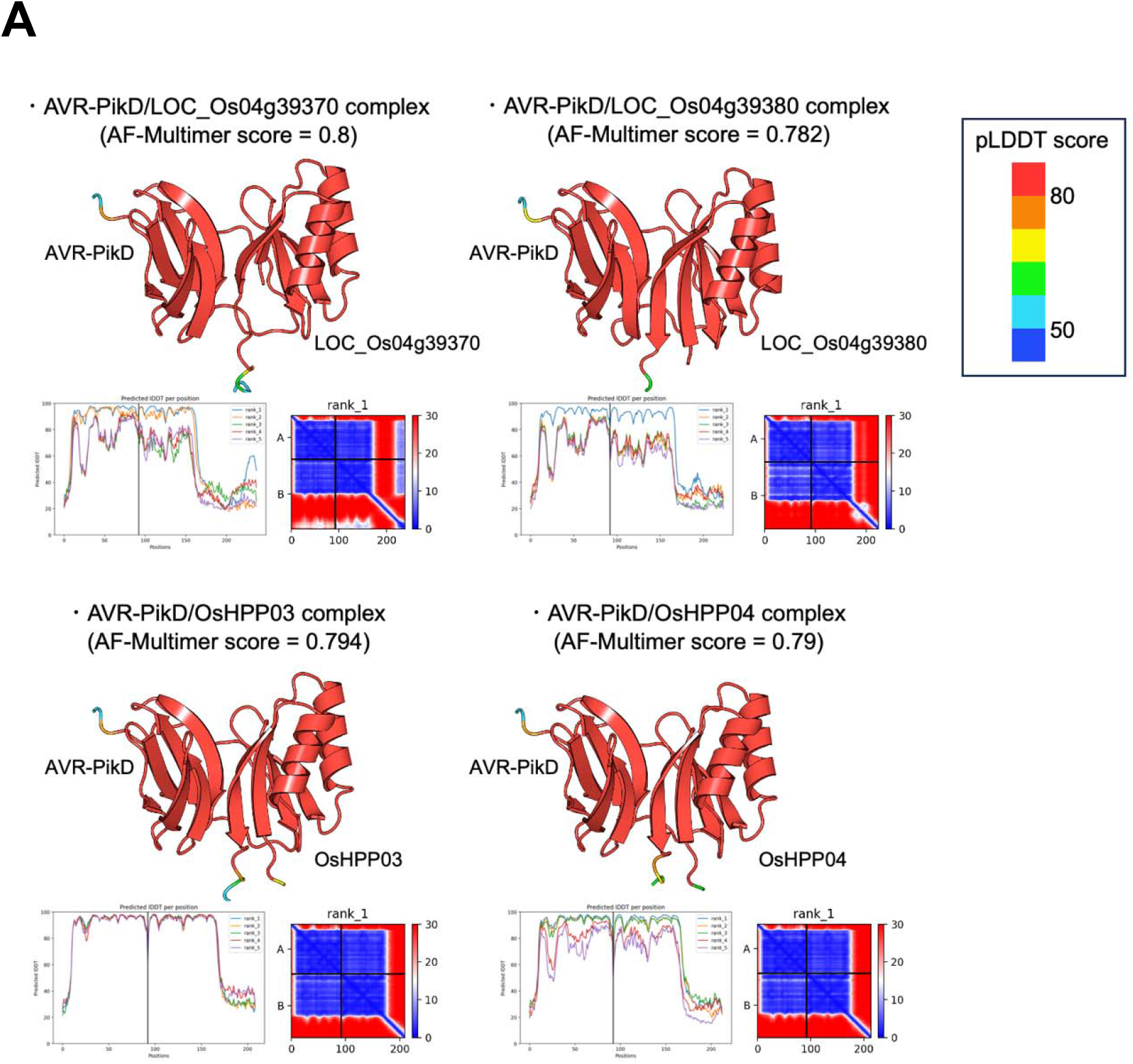
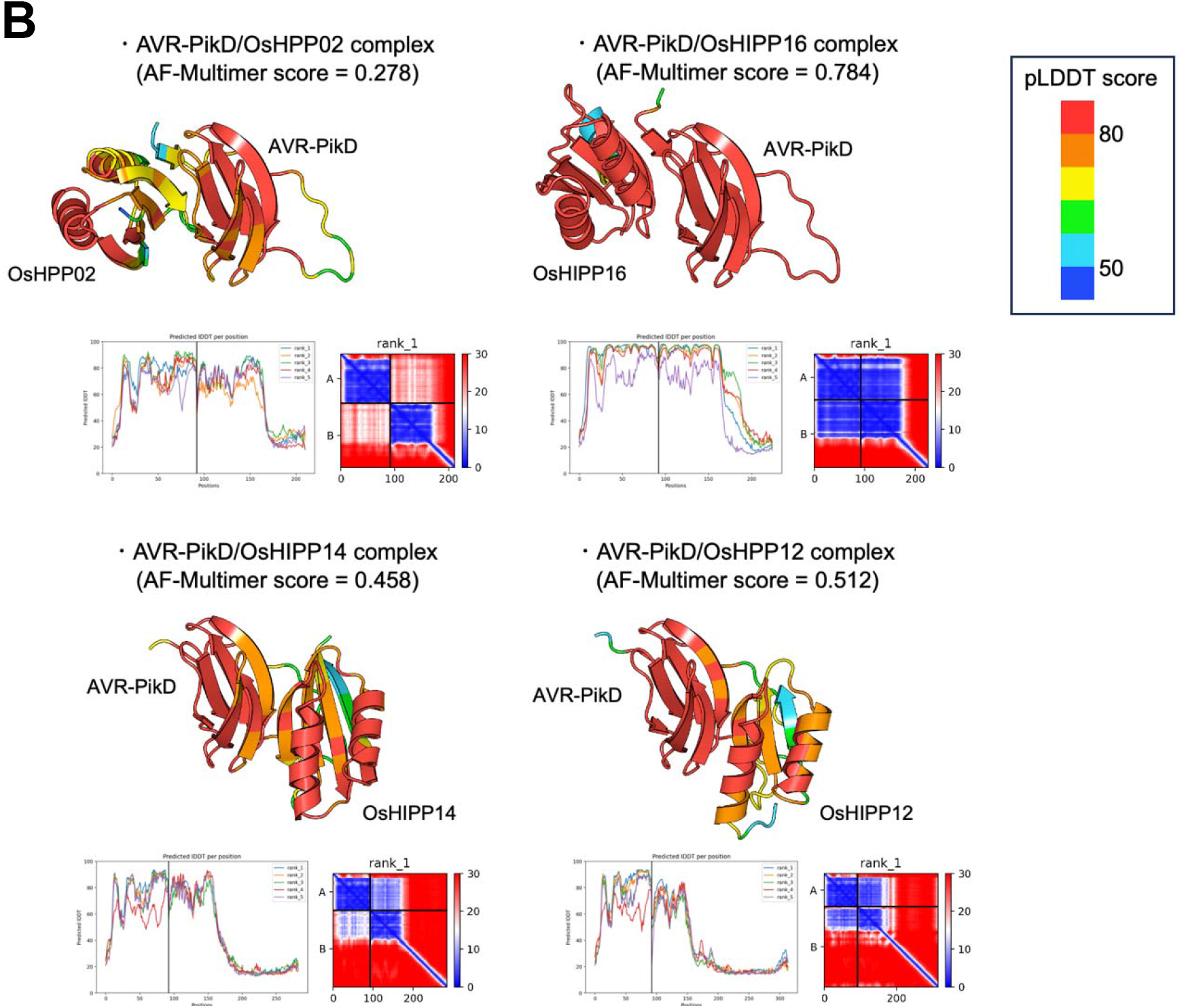
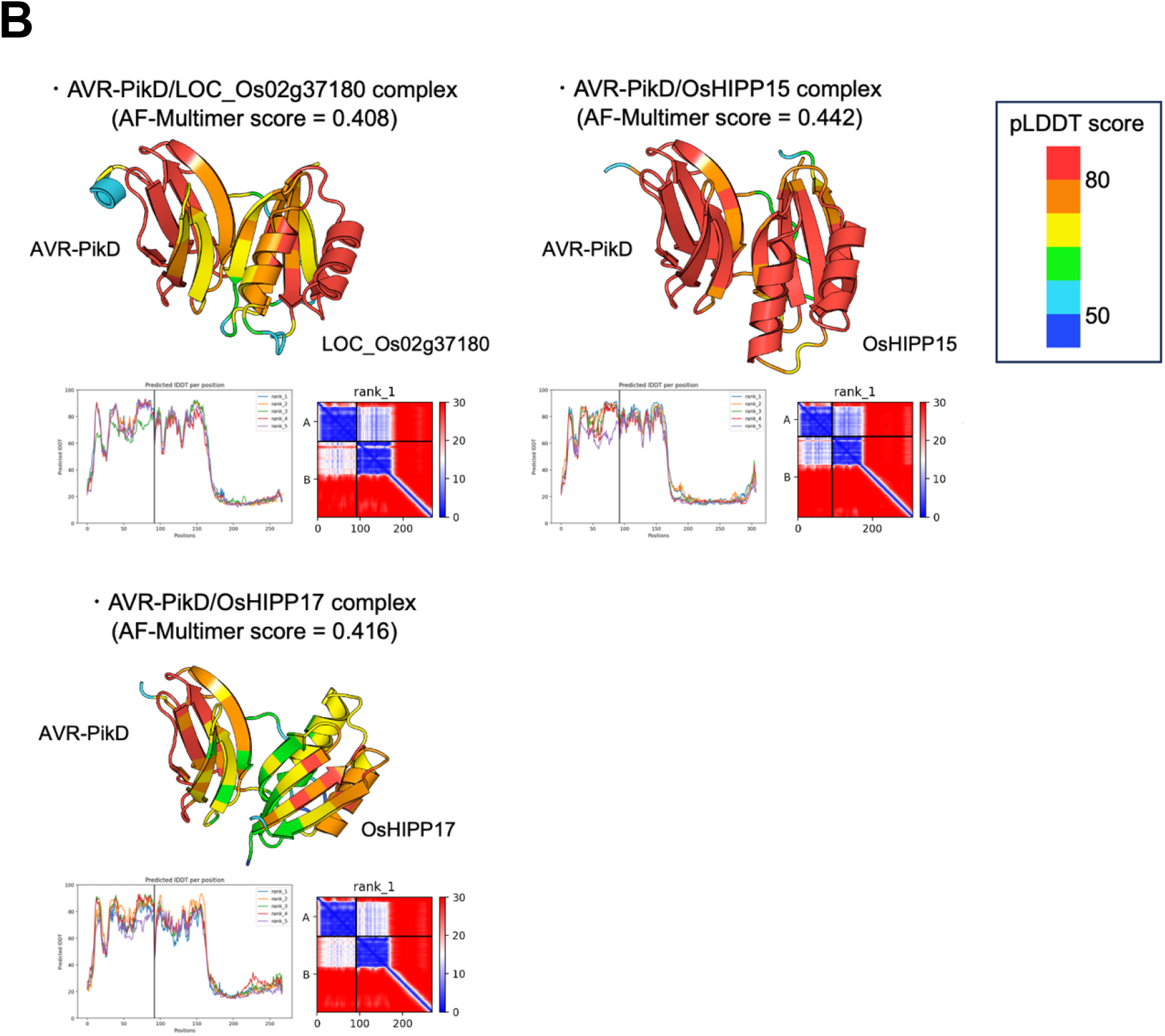
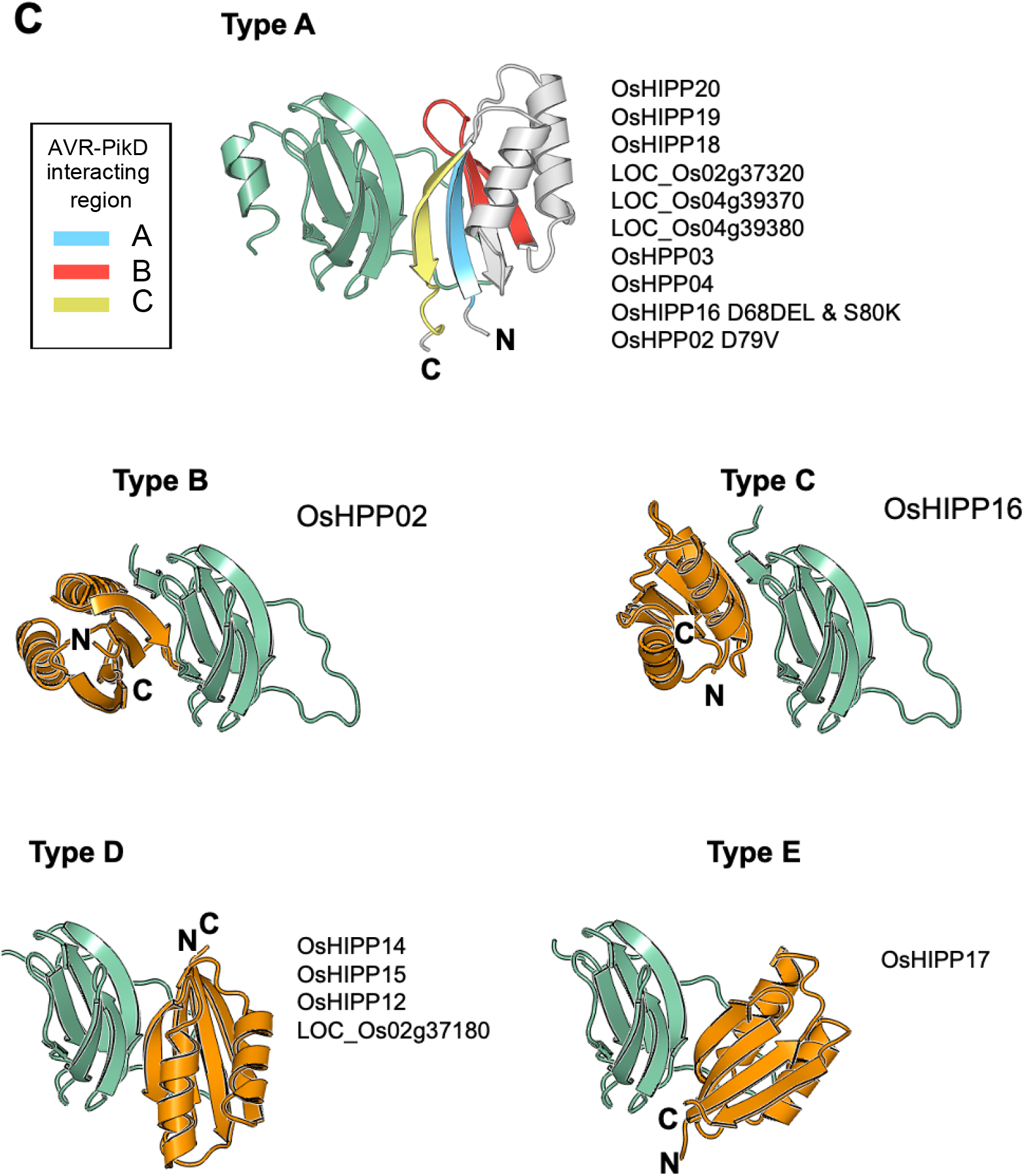
**A:** AVR-PikD/sHMA (OsHIPP20, OsHIPP19, OsHIPP18 and LOC_Os02g37320) complex predictions were generated in ColabFold_v1.5.2. For each protein, predicted binding structure (top), pLDDT (bottom left), predicted aligned error (bottom right) are shown. Predicted regions with low pLDDT score (< 50) are not displayed. AlphaFold (AF) -multimer score = 0.8*ipTM + 0.2*pTM (Yin et al. 2022). **A:** AVR-PikD/sHMA (LOC_Os04g39370, LOC_Os04g39380, OsHPP03 and OsHPP04) complex predictions were generated in ColabFold_v1.5.2. For each protein, predicted binding structure (top), pLDDT (bottom left), predicted aligned error (bottom right) are shown. Predicted regions with low pLDDT score (< 50) are not displayed. AlphaFold (AF) -multimer score = 0.8*ipTM + 0.2*pTM (Yin et al. 2022). **B:** AVR-PikD/sHMA (OsHPP02, OsHIPP16, OsHIPP14 and OsHIPP12) complex predictions were generated in ColabFold_v1.5.2. For each protein, predicted binding structure (top), pLDDT (bottom left), predicted aligned error (bottom right) are shown. Predicted regions with low pLDDT score (< 50) are not displayed. AlphaFold (AF) -multimer score = 0.8*ipTM + 0.2*pTM (Yin et al. 2022). **B:** AVR-PikD/sHMA (LOC_Os02g37180, OsHIPP15 and OsHIPP17) complex predictions were generated in ColabFold_v1.5.2. For each protein, predicted binding structure (top), pLDDT (bottom left), predicted aligned error (bottom right) are shown. Predicted regions with low pLDDT score (< 50) are not displayed. AlphaFold (AF) -multimer score = 0.8*ipTM + 0.2*pTM (Yin et al. 2022). **C:** Five types of AVR-PikD (light green) / sHMA (white and orange) complexes (Type A to E) predicted by ColabFold. Predicted complex structure of AVR-PikD and sHMA that were shown to interact in Y2H (**Fig.1**) all belonged to Type A, while those of AVR-PikD and sHMA non-interacting in Y2H belonged to either of Type B, C, D, E.

**Fig. S6.**
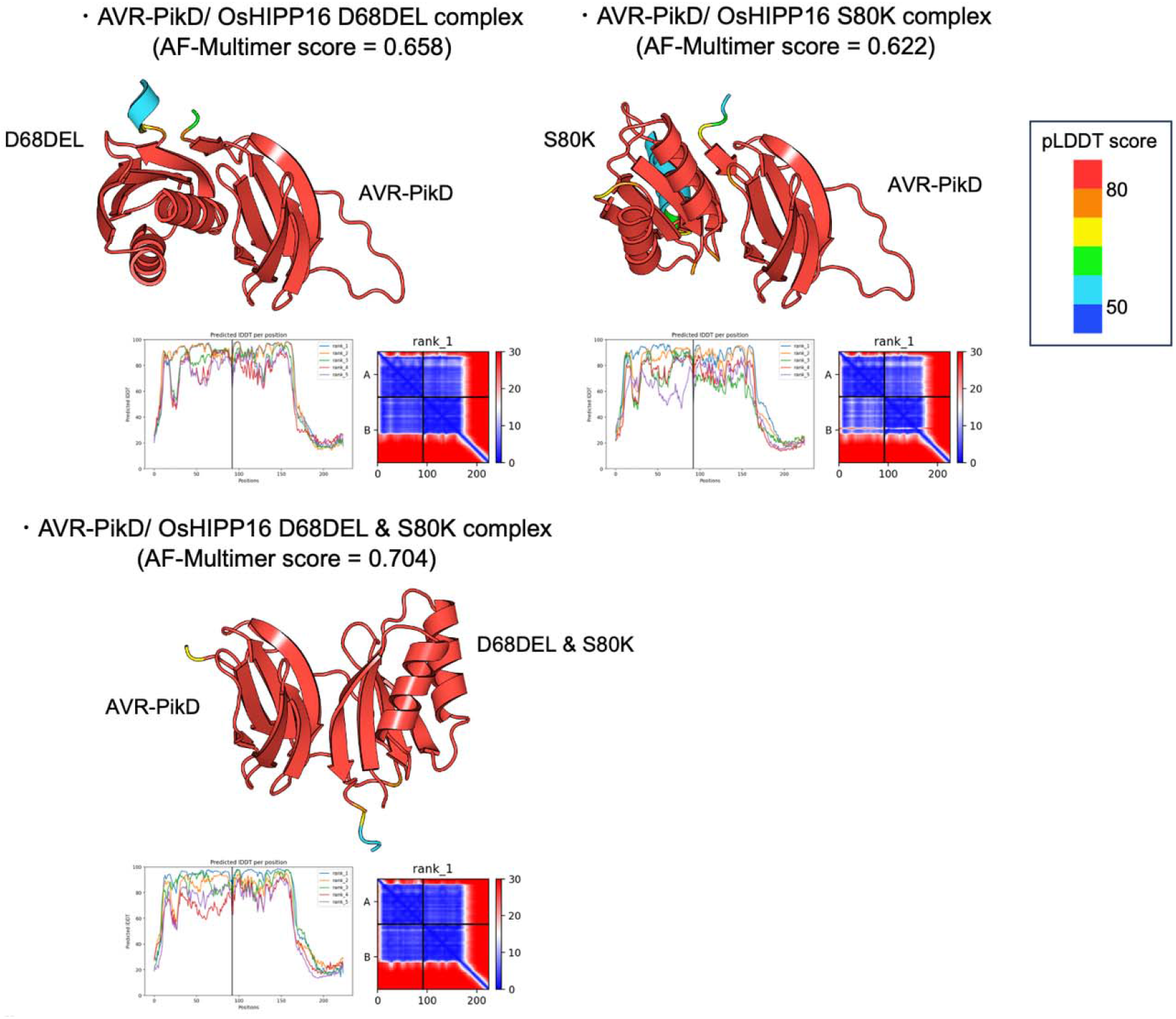
**A:** AVR-PikD/sHMA (OsHIPP16_D68DEL, OsHIPP16_S80K, OsHIPP16_D68DEL_S80K) complex predictions were generated in ColabFold_v1.5.2. For each protein, predicted binding structure (top), pLDDT (bottom left), predicted aligned error (bottom right) are shown. Predicted regions with low pLDDT score (< 50) are not displayed. AlphaFold (AF) -multimer score = 0.8*ipTM + 0.2*pTM (Yin et al. 2022).

**Fig. S7.**
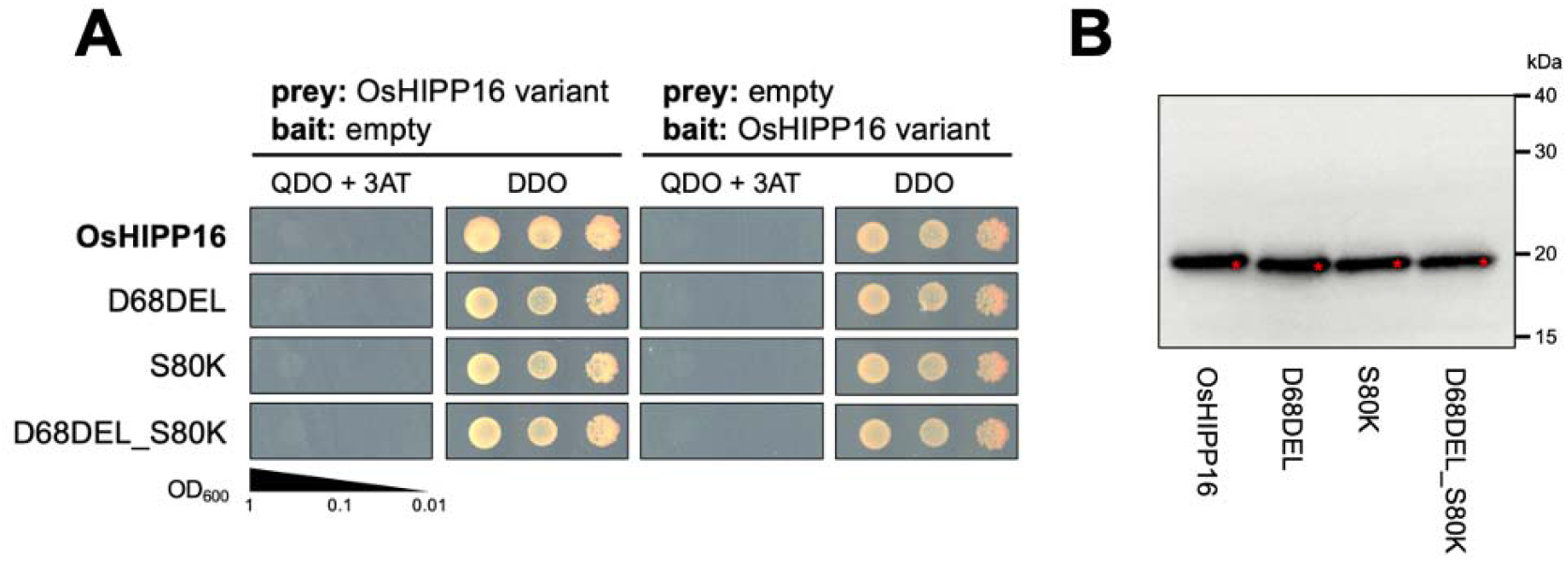
**A:** Y2H interactions between the variants of OsHIPP16 (OsHIPP16, OsHIPP16_D68DEL, OsHIPP16_S80K, OsHIPP16_D68DEL_S80K) and the empty vector products. **B:** Western blot analysis confirms protein production in the AlphaScreen as shown in Figure 2D. The sHMA proteins were tagged with the FLAG epitope and detected by anti-FLAG antibody. The protein bands expressed from vectors are marked by red asterisks. The positions of molecular size marker are indicated on the right (kDa).

**Fig. S8.**
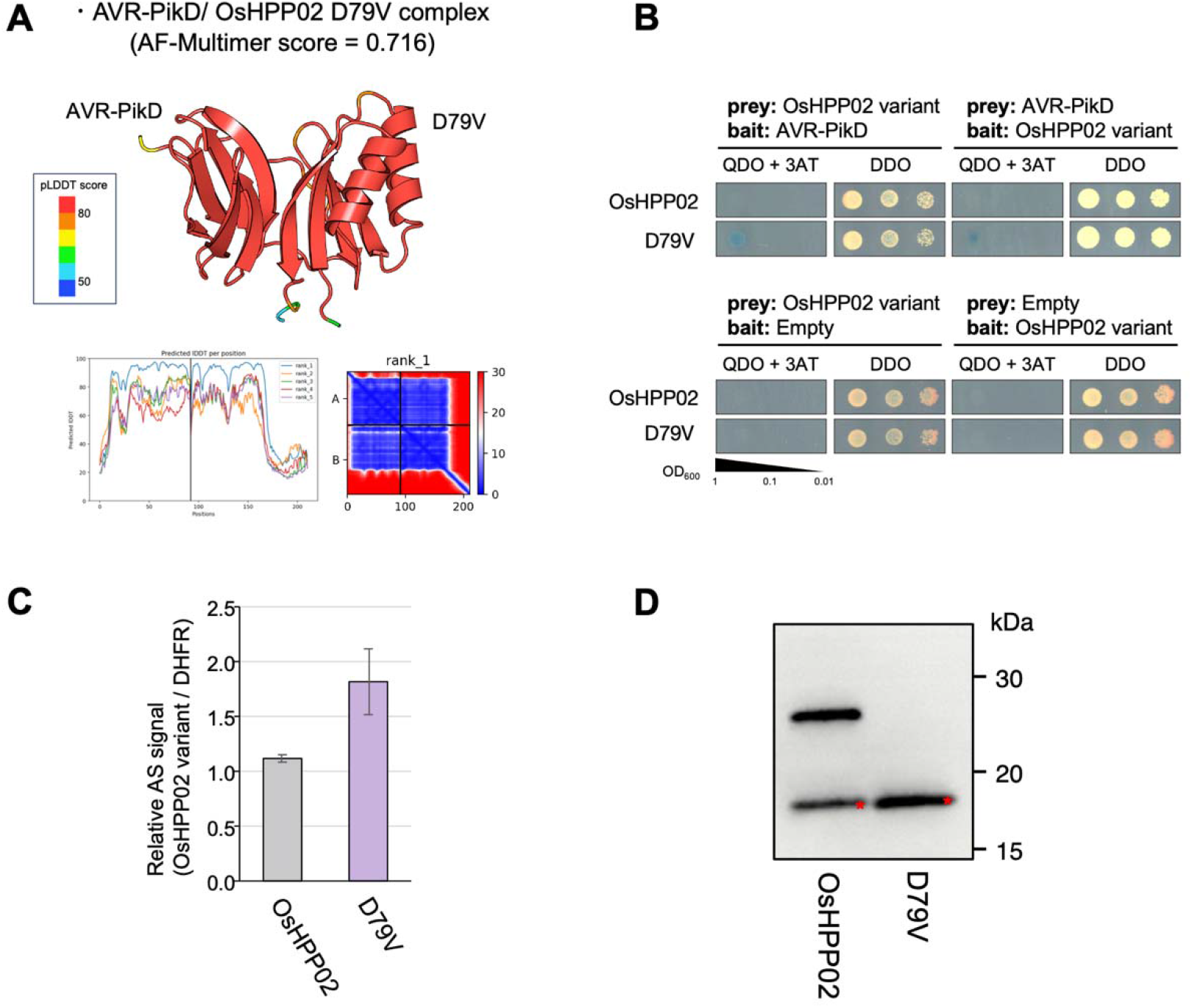
A mutant version of OsHPP02 binds AVR-PikD. **A:** AVR-PikD/sHMA (OsHPP02_D79V) complex predictions were generated in ColabFold_v1.5.2. Predicted binding structure (top), pLDDT (bottom left), predicted aligned error (bottom right) are shown. Predicted regions with low pLDDT score (< 50) are not displayed. AlphaFold (AF) -multimer score = 0.8*ipTM + 0.2*pTM (Yin et al. 2022). **B:** Y2H interaction assay between AVR-PikD and OsHPP02 and OsHPP02_D79V. **C:** AlphaScreen interaction assay between AVR-PikD and OsHPP02 and OsHPP02_D79V. Relative signal strength as compared to that between OsHIPP02 and DHFR (negative control) is given. **D:** Western blot analysis confirms protein production in the AlphaScreen as shown in **C**. The sHMA proteins were tagged with the FLAG epitope and detected by anti-FLAG antibody. The protein bands expressed from vector are marked by red asterisks. The positions of molecular size marker are indicated on the right (kDa).

**Fig. S9.**
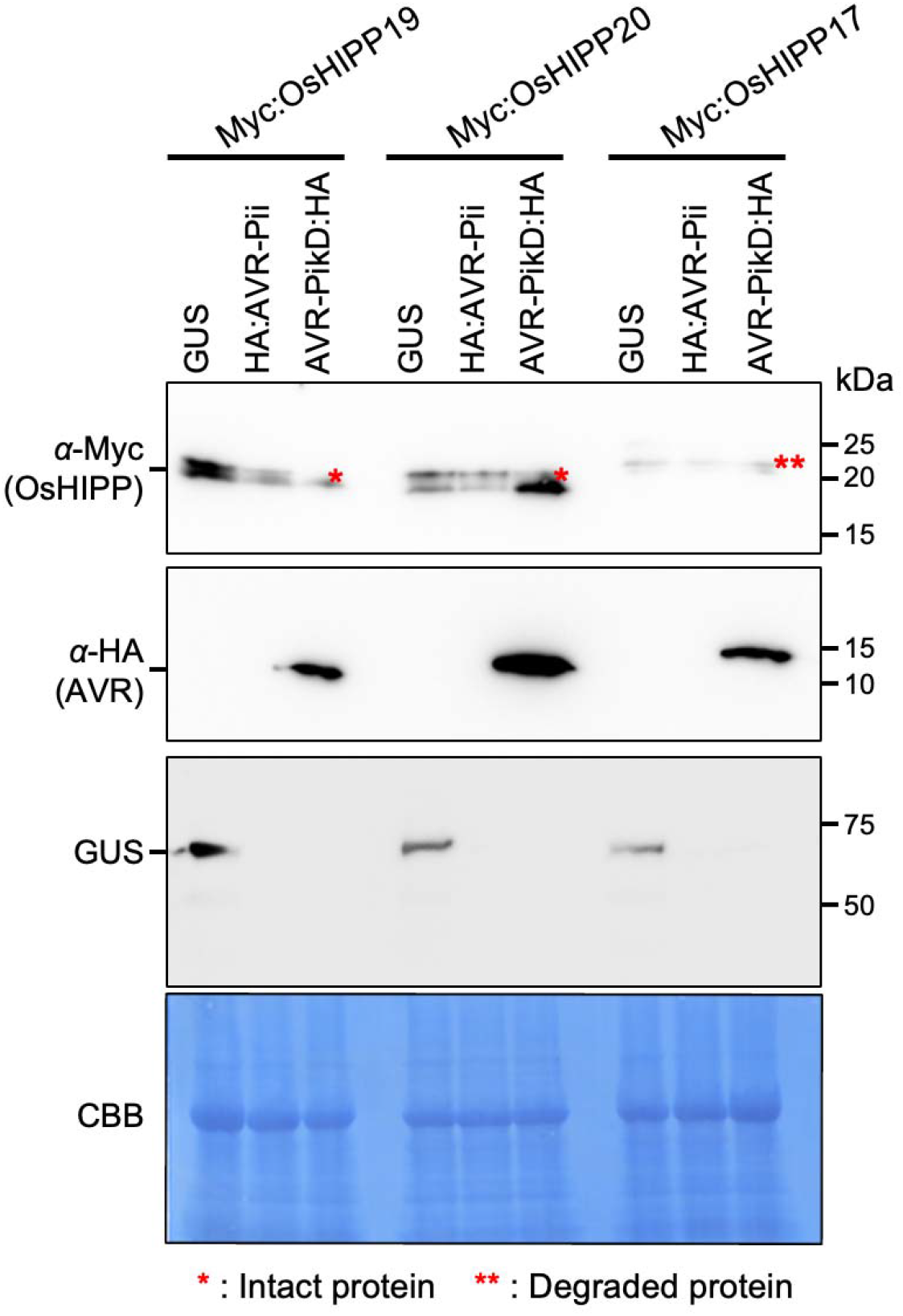
AVR-PikD stabilizes sHMAs in planta. The results for pellet fraction after fractionation of leaf extract are shown. AVR-PikD seems to accumulate in the pellet fraction when it does not bind sHMA.

**Fig. S10.**
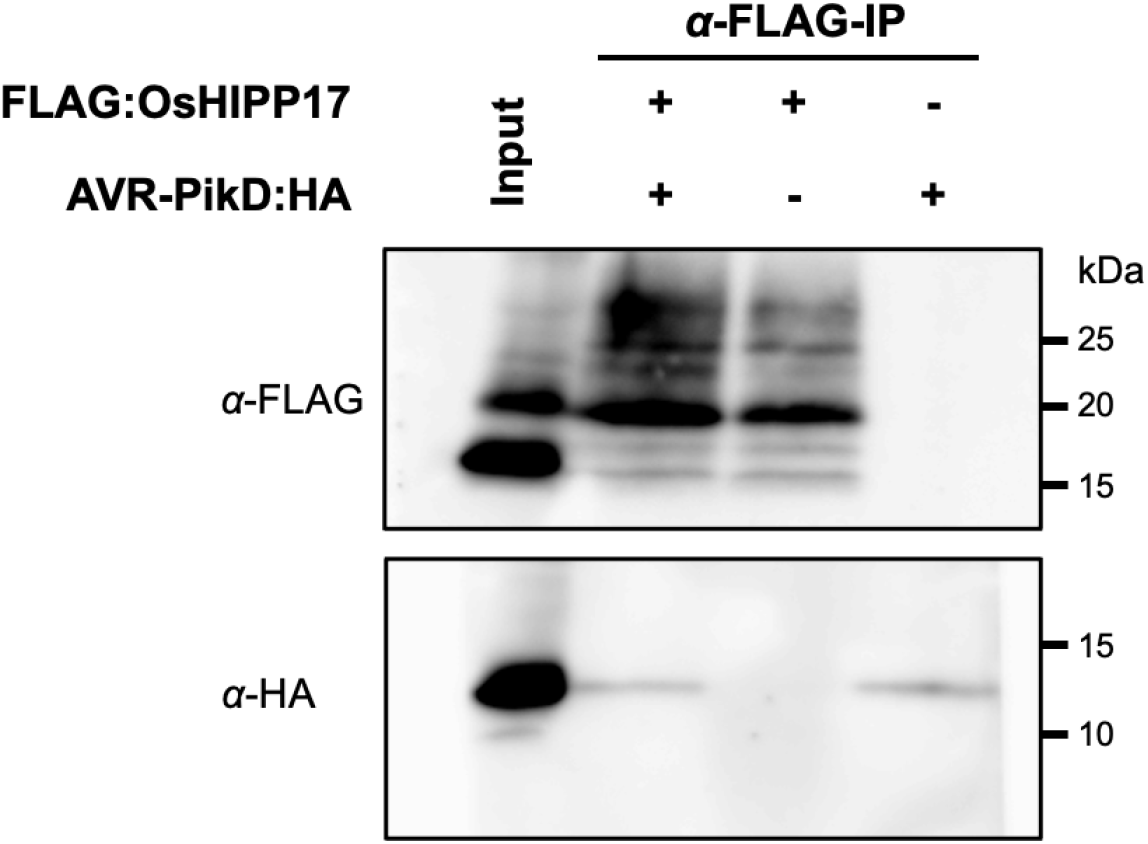
Co-immunoprecipitation experiment shows AVR-PikD does not bind OsHIPP17. Binding assay between OsHIPP17 and AVR-PikD. Epitope-tagged proteins, AVR-PikD:HA and FLAG:OsHIPP17 were expressed in *Nicotiana benthamiana* leaves. The leaf extract was applied to an anti- FLAG antibody column and the bound proteins were detected by an anti-FLAG antibody (top) and an anti- HA antibody (bottom). AVR-PikD:HA band detected in the anti-HA blot after co-immunoprecipitation is caused by non-specific weak binding of AVR-PikD:HA to anti-FLAG antibody column.

**Fig. S11 (Supplement to Fig. 3B).**
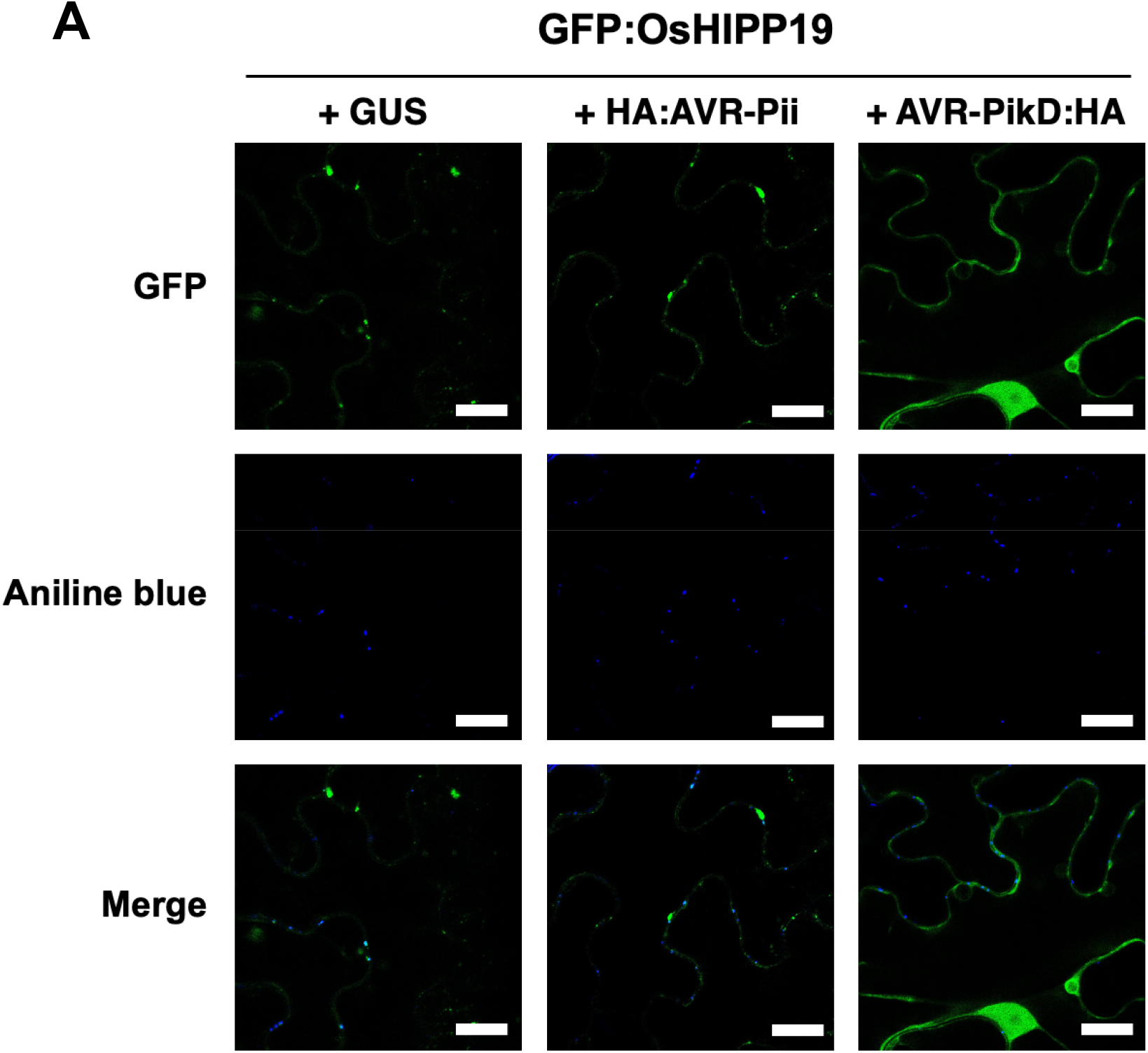
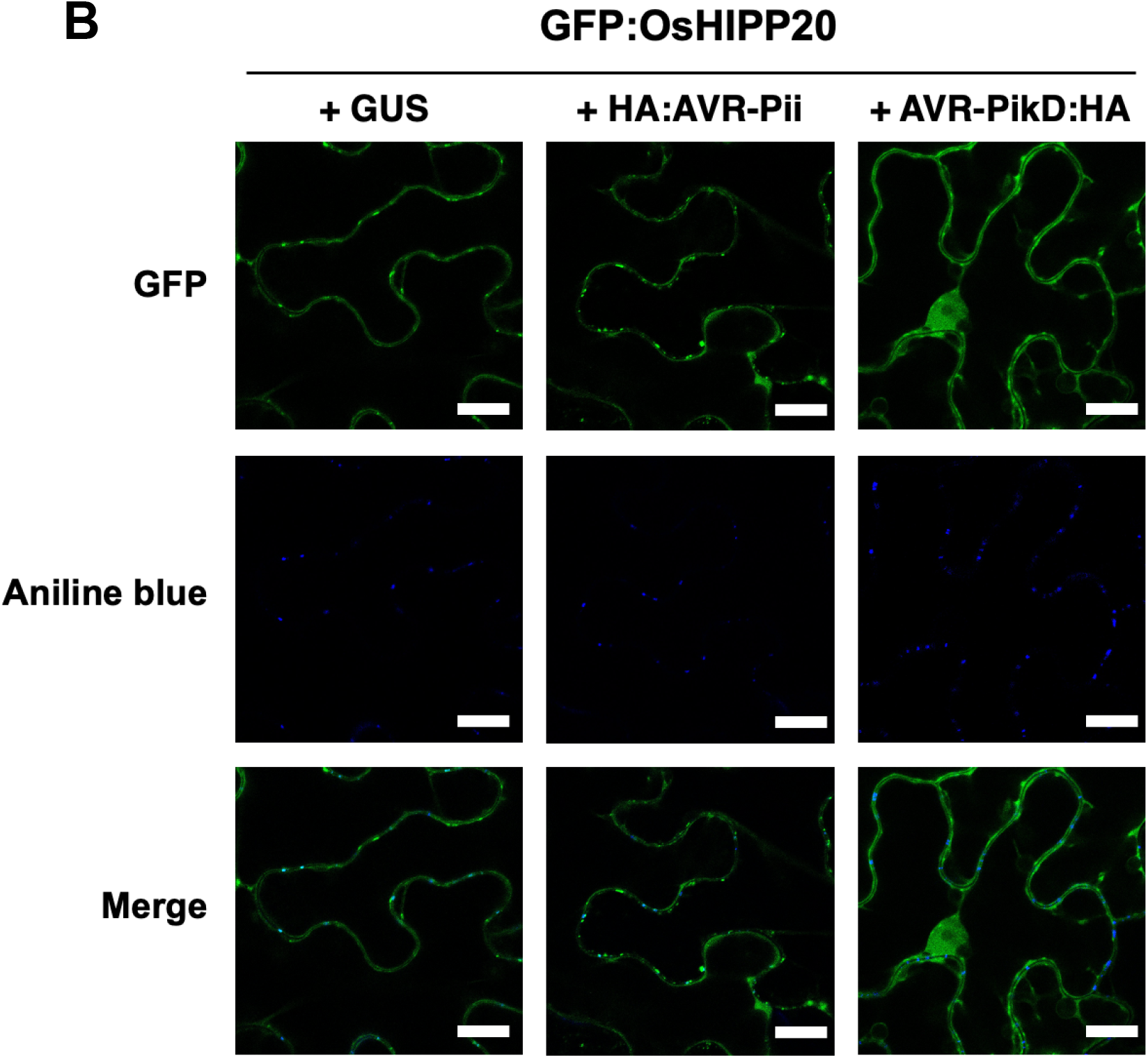
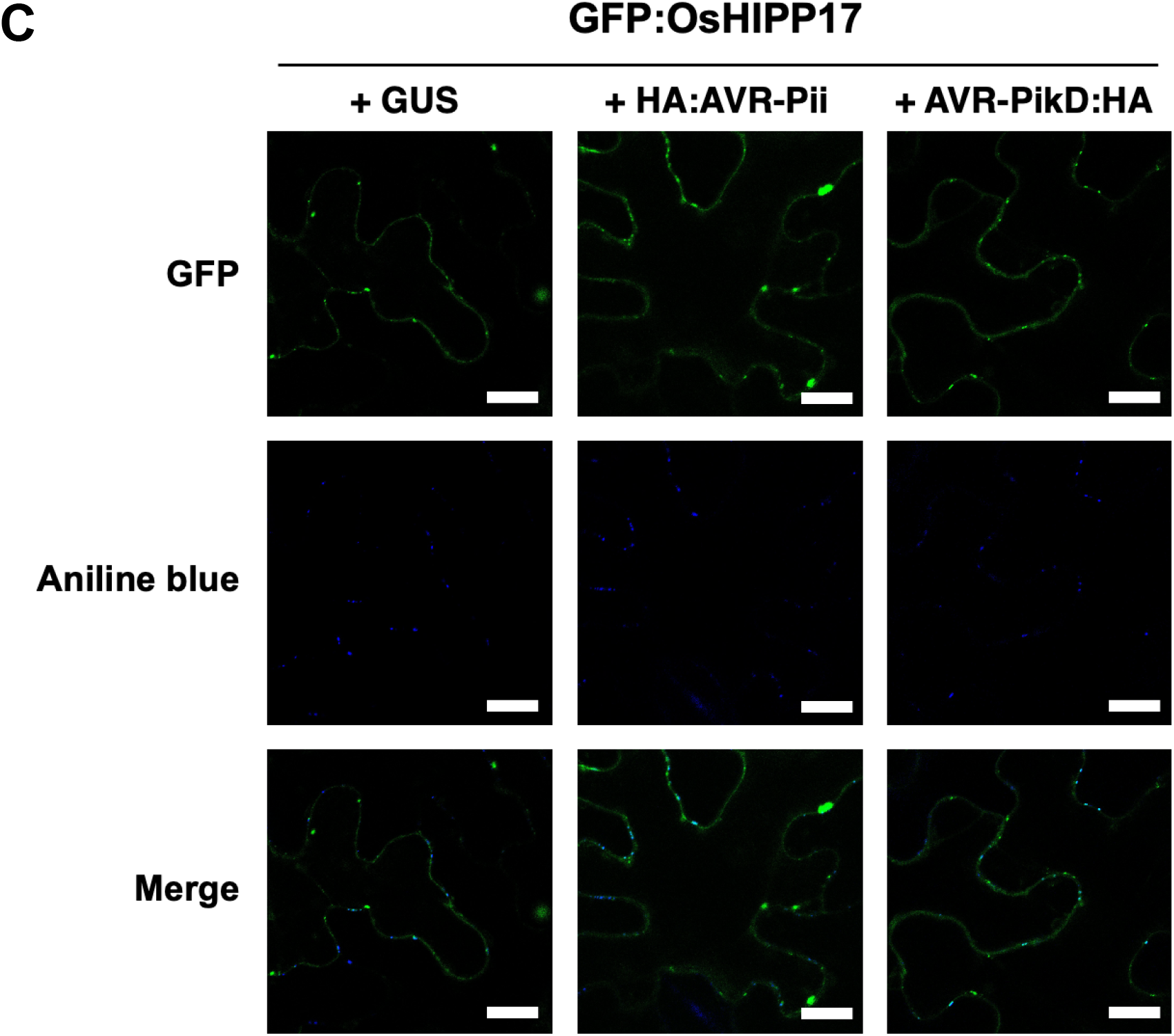
**A:** Subcellular localization of GFP:OsHIPP19 expressed in *N. benthamiana* leaves in the presence of GUS, HA:AVR-Pii and AVR-PikD:HA. Scale bar: 20 μm. **B:** Subcellular localization of GFP:OsHIPP20 expressed in *N. benthamiana* leaves in the presence of GUS, HA:AVR-Pii and AVR-PikD:HA. Scale bar: 20 μm. **C:** Subcellular localization of GFP:OsHIPP17 expressed in *N. benthamiana* leaves in the presence of GUS, HA:AVR-Pii and AVR-PikD:HA. Scale bar: 20 μm.

**Fig. S12.**
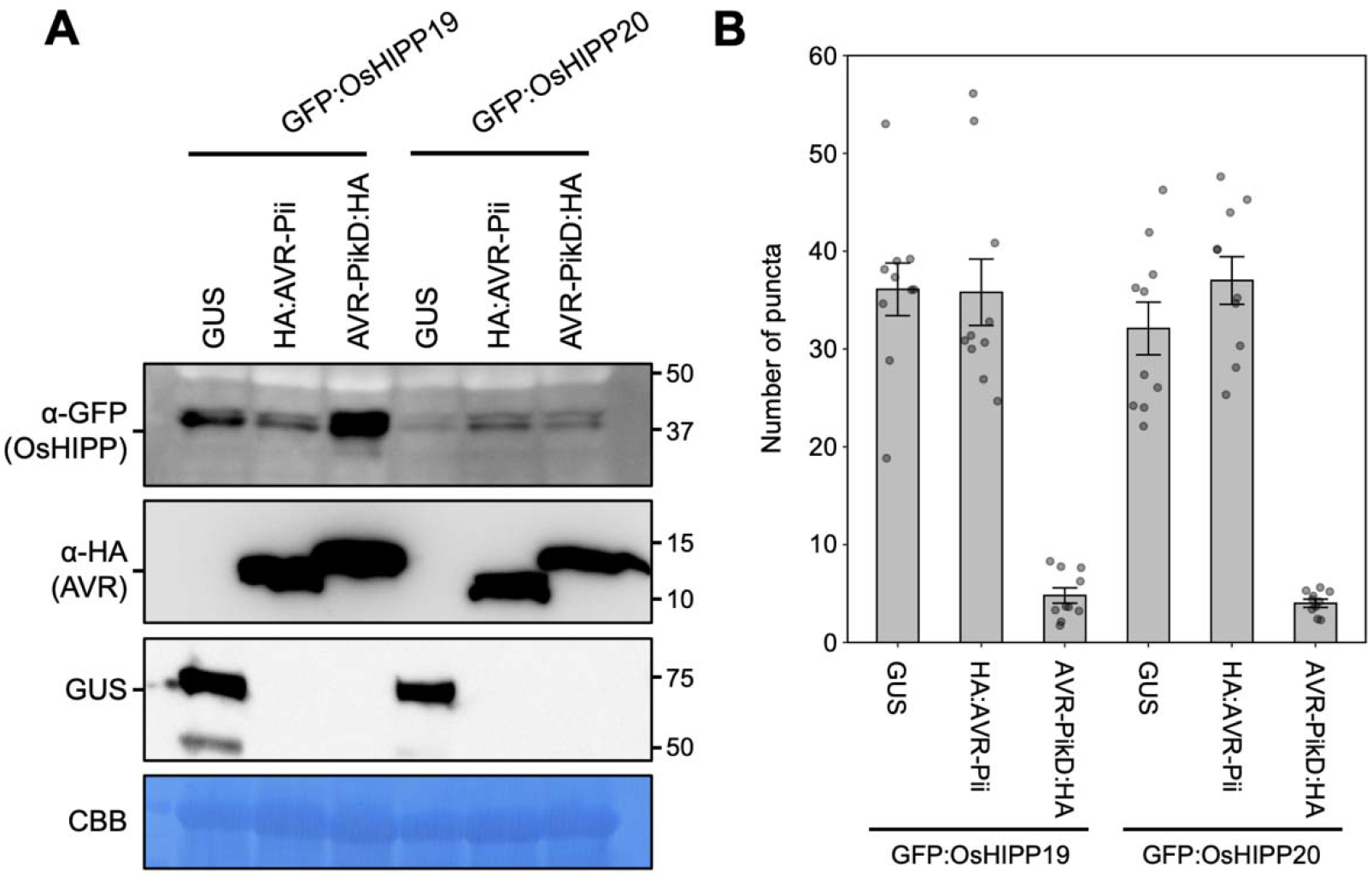
AVR-Pik-D binding affects subcellular localization of sHMAs expressed in *N. benthamiana*. **A:** Results of Western blot analysis of proteins expressed in *N. benthamiana* leaves as shown in **Fig. 3C**. **B:** Histograms showing the number of GFP:OsHIPP19 and GFP:OsHIPP20 puncta structure in *N. benthamiana* cells in the presence of GUS, HA:AVR-Pii and AVR-PikD:HA.

**Fig. S13.**
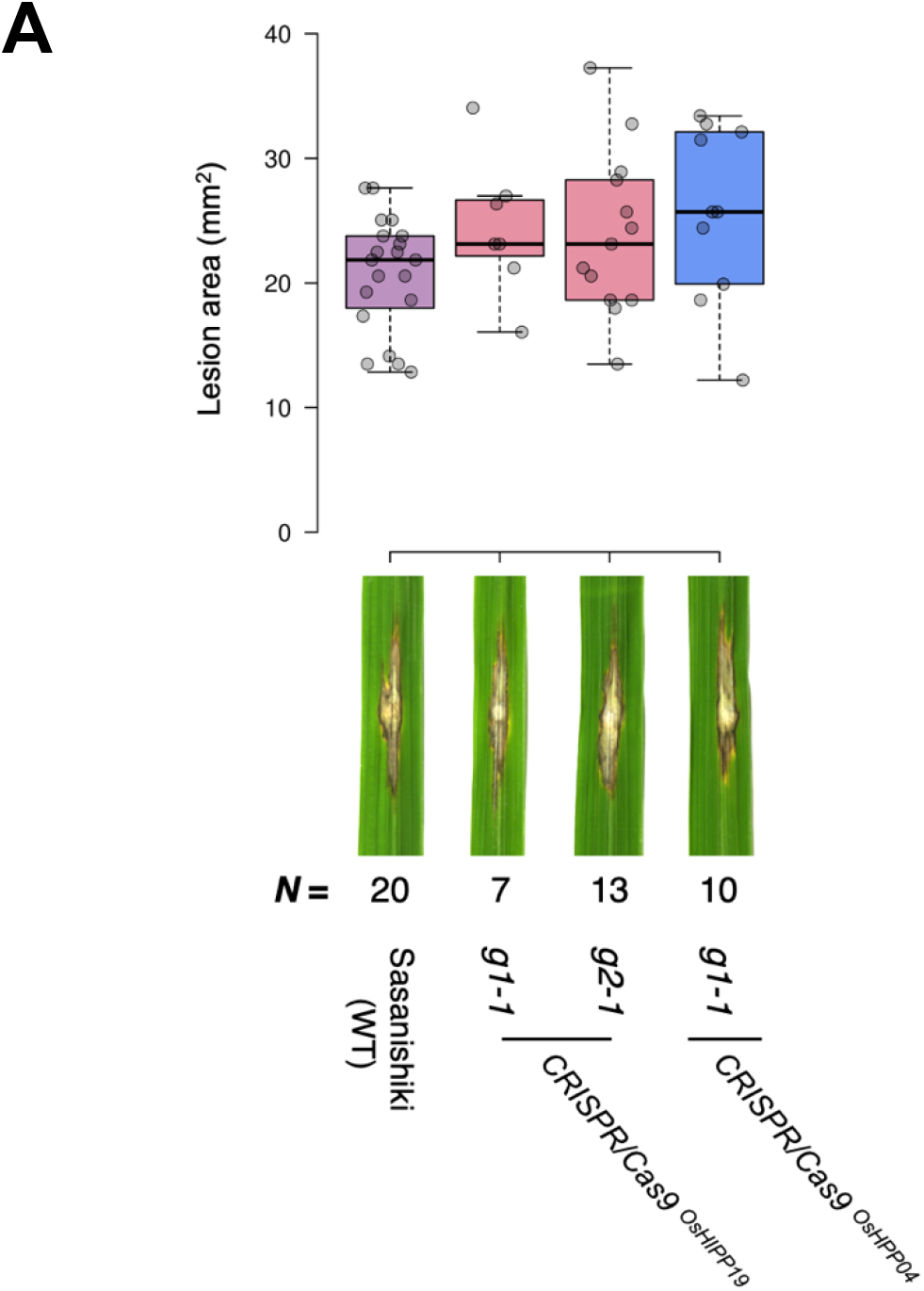
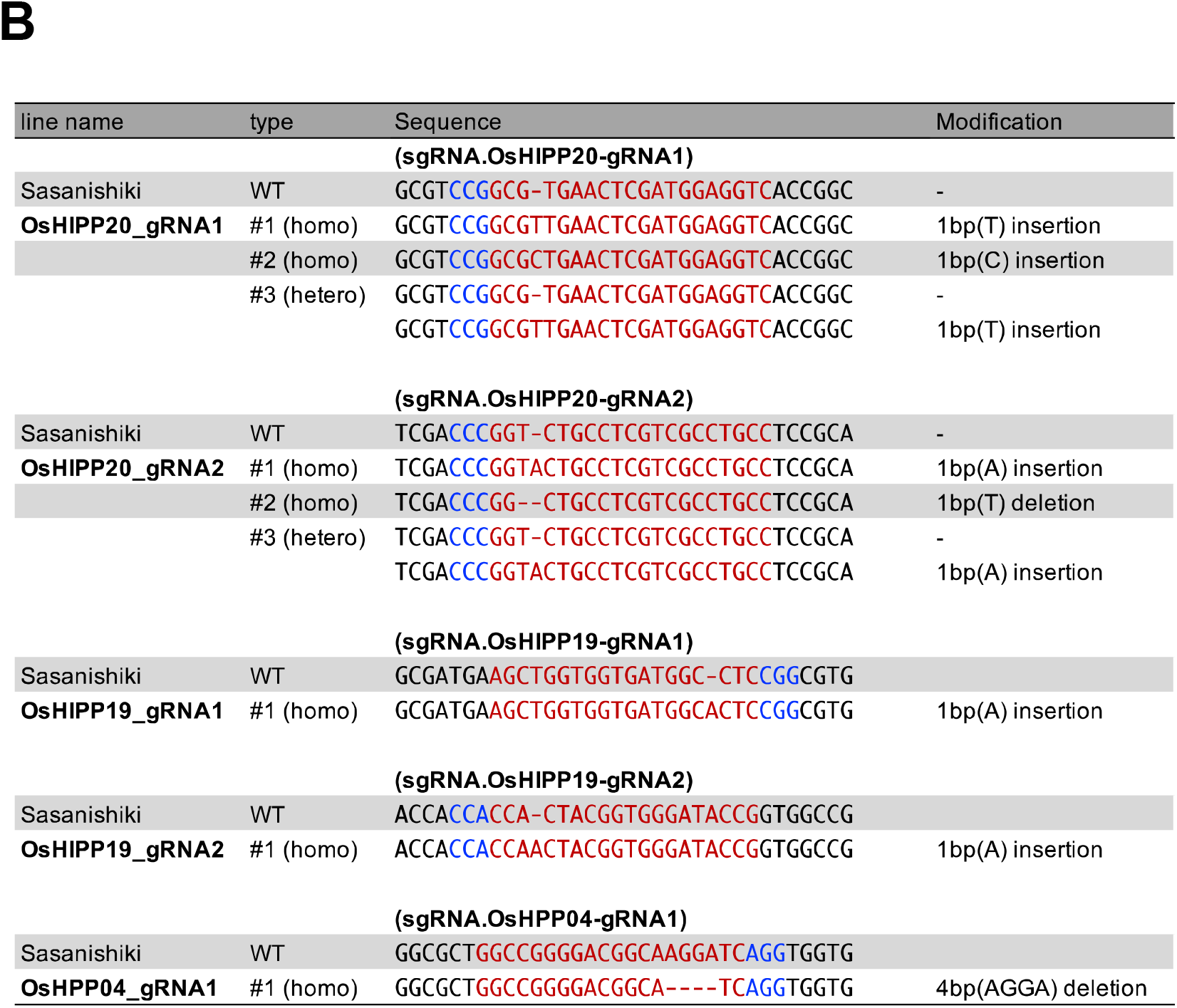
Knockout of *OsHIPP19* and *OsHPP04* does not alter susceptibility of rice against a compatible *M. oryzae* isolate. **A:** A compatible *M. oryzae* isolate Sasa2 was punch inoculated onto the leaves of rice cultivar Sasanishiki as well as the sHMA-knockout lines of Sasanishiki (oshipp19#1, oshipp19#2 and oshpp04). Box plots show lesion area sizes in the rice lines (top). Photos of typical lesions developed on the leaves after inoculation of *M. oryzae* (bottom). The number of leaves used for experiments are indicated below. **B:** A table showing guide RNA and transgenic line nomenclature (left), location of guide RNA used for CRISPR/Cas9 mutagenesis (center) and the resulting nucleotide changes (center and right). PAM is indicated with blue and the sgRNA sequence is indicated with red letters.

**Table S2.**
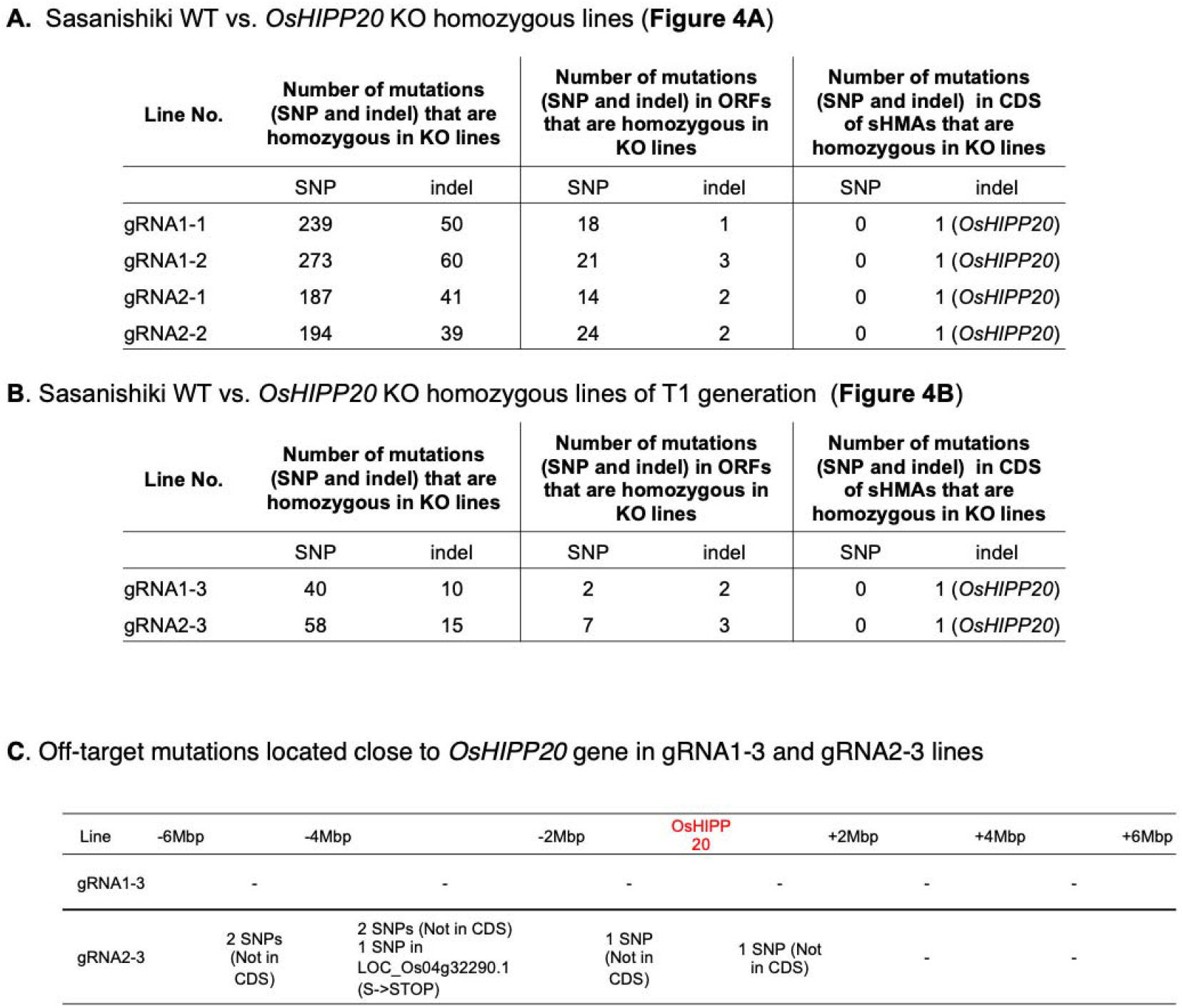
Summary of whole genome sequence comparison between the wild type Sasanishiki (WT) and *OsHIPP20* KO lines.

**Fig. S14.**
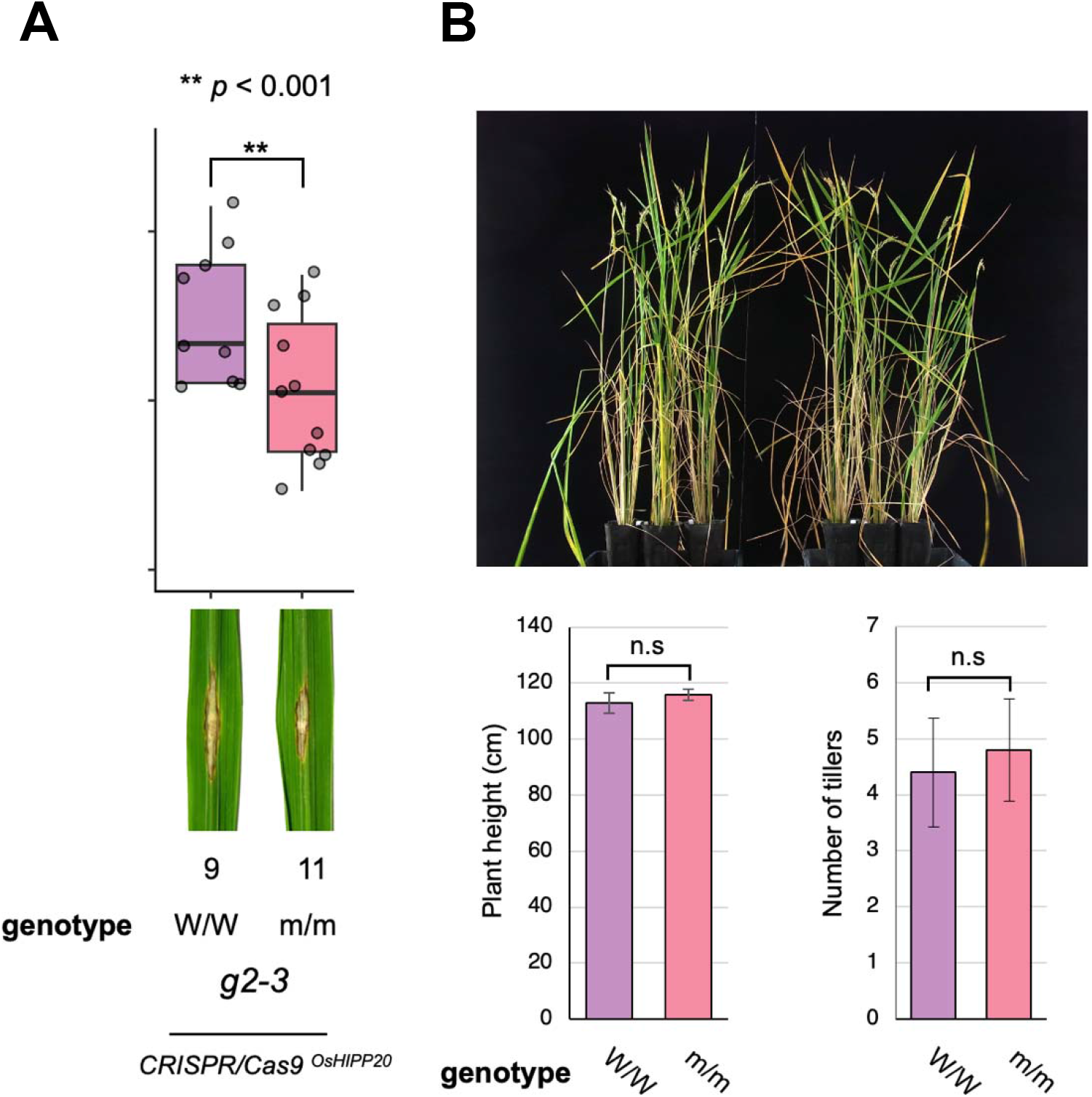
A Sasanishiki line (g2-3) in which both *OsHIPP20* and *LOC_Os04g32290.1* gene were knocked out shows enhanced resistance against a compatible *M. oryzae* but does not show growth defect. A: Punch inoculation results of T2 generation of homozygous KO rice plants. **B:** No growth defect in *OsHIPP20* + *LOC_Os04g32290.1* KO line as compared to the wild type. Top: Overview of the W/W and m/m plants. Bottom: Box plot showing plant height (left) and tiller number distribution of W/W and m/m plants.

**Dataset S1.**
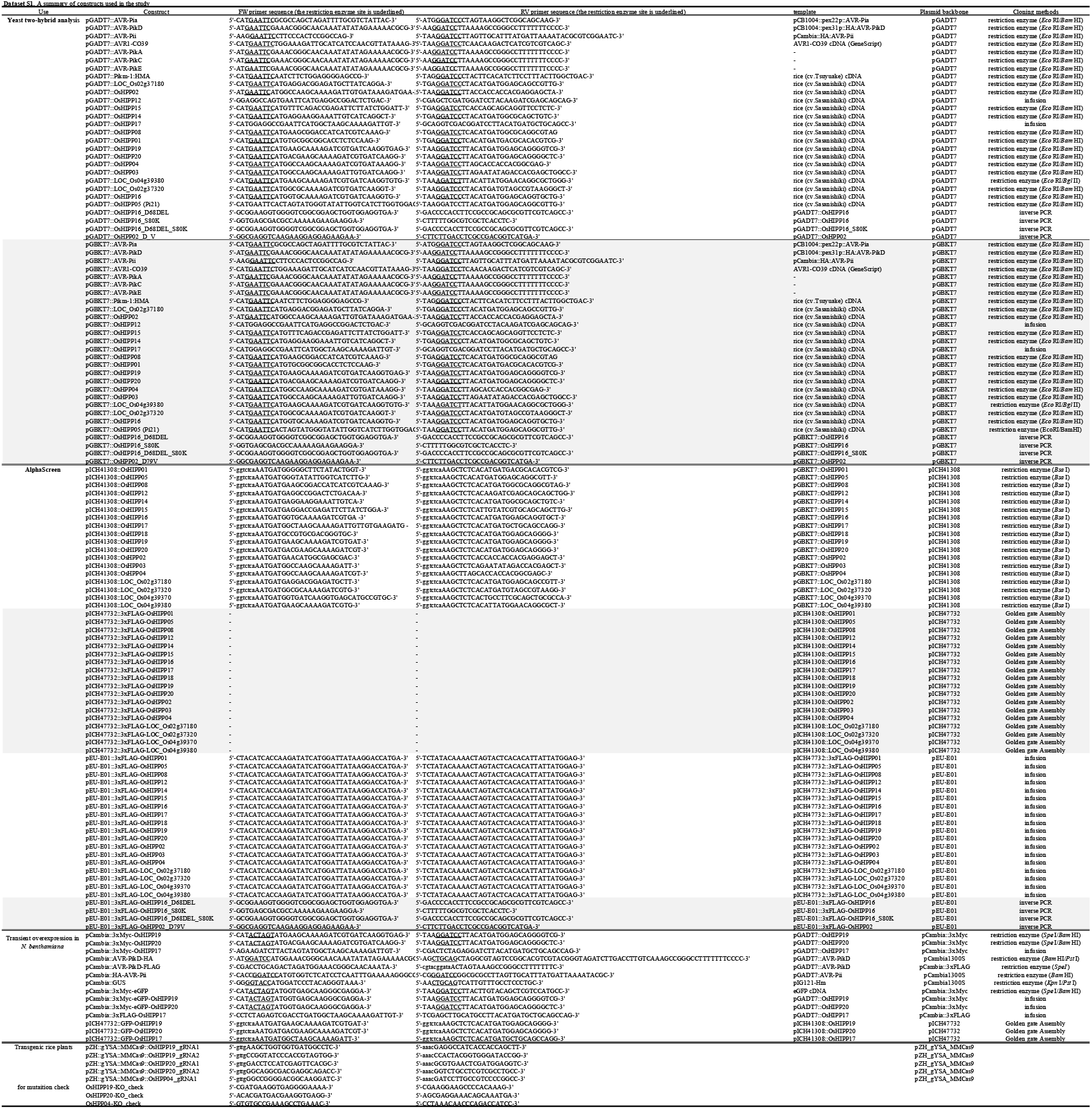
A summary of constructs used in the study.

## References

1. S. A. Hogenhout, R. A. Van der Hoorn, R. Terauchi, S. Kamoun, Emerging concepts in effector biology of plant-associated organisms. Mol. Plant-Microbe Interact. MPMI 22, 115–122 (2009).

2. S. Cesari, et al., A novel conserved mechanism for plant NLR protein pairs: the “integrated decoy” hypothesis. Front. Plant Sci. 5, 606 (2014).

3. C.H. Wu, et al., The “sensor domains” of plant NLR proteins: more than decoys? Front. Plant Sci. 6, 134 (2015).

4. P. F. Sarris, et al., Comparative analysis of plant immune receptor architectures uncovers host proteins likely targeted by pathogens. BMC Biol. 14, 8 (2016).

5. T. Kroj, et al., Integration of decoy domains derived from protein targets of pathogen effectors into plant immune receptors is widespread. New Phytol. 210, 618–626 (2016).

6. E. Baggs, G. Dagdas, K. V. Krasileva, NLR diversity, helpers and integrated domains: making sense of the NLR IDentity. Curr. Opin. Plant Biol. 38, 59–67 (2017).

7. P. C. Bailey, et al., Dominant integration locus drives continuous diversification of plant immune receptors with exogenous domain fusions. Genome Biol. 19, 23 (2018).

8. Y. Okuyama, et al., A multifaceted genomics approach allows the isolation of the rice Pia-blast resistance gene consisting of two adjacent NBS-LRR protein genes. Plant J. 66, 467–479 (2011).

9. I. Ashikawa, et al., Two adjacent nucleotide-binding site-leucine-rich repeat class genes are required to confer Pikm-specific rice blast resistance. Genetics. 180, 2267–2276 (2008).

10. K. Yoshida, et al., Association genetics reveals three novel avirulence genes from the rice blast fungal pathogen *Magnaporthe oryzae*. Plant Cell. 21, 1573–1591 (2009).

11. K. De Guillen, et al., Structure Analysis Uncovers a Highly Diverse but Structurally Conserved Effector Family in Phytopathogenic Fungi. PLoS Pathog. 11, e1005228 (2015).

12. A. Maqbool, et al., Structural basis of pathogen recognition by an integrated HMA domain in a plant NLR immune receptor. Elife. 4, e08709 (2015).

13. H. Kanzaki, et al., Arms race co-evolution of *Magnaporthe oryzae AVR-Pik* and rice *Pik* genes driven by their physical interactions. Plant J. 72, 894–907 (2012).

14. S. Cesari, et al., The rice resistance protein pair RGA4/RGA5 recognizes the *Magnaporthe oryzae* effectors AVR-Pia and AVR1-CO39 by direct binding. Plant Cell. 25, 1463–1481 (2013).

15. J. C. De la Concepcion, et al., Polymorphic residues in rice NLRs expand binding and response to effectors of the blast pathogen. Nat. Plants. 4, 576–585 (2018).

16. A. Białas, et al., Lessons in Effector and NLR Biology of Plant-Microbe Systems. Mol. Plant-Microbe Interact. MPMI 31, 34–45 (2018).

17. J. C. De la Concepcion, et al., Protein engineering expands the effector recognition profile of a rice NLR immune receptor. Elife. 8, e47713 (2019).

18. J. H. R. Maidment, et al., Effector target-guided engineering of an integrated domain expands the disease resistance profile of a rice NLR immune receptor. Elife. 12, e81123 (2023).

19. L. Banci, et al., Solution structure of the yeast copper transporter domain Ccc2a in the apo and Cu(I)-loaded states. J. Biol. Chem. 276, 8415–8426 (2001).

20. L. Guo, et al., Specific recognition of two MAX effectors by integrated HMA domains in plant immune receptors involves distinct binding surfaces. Proc. Natl. Acad. Sci. U. S. A. 115, 11637– 11642 (2018).

21. J. B. De Abreu-Neto, et al., Heavy metal-associated isoprenylated plant protein (HIPP): characterization of a family of proteins exclusive to plants. FEBS J. 280, 1604–1616 (2013).

22. S. Fukuoka, et al., Loss of function of a proline-containing protein confers durable disease resistance in rice. Science. 325, 998–1001 (2009).

23. N.N. Shi, et al., Virulence structure of *Magnaporthe oryzae* populations from Fujian Province, China. Canadian J. Plant Pathol. 40, 542–550 (2018).

24. Y. Kawahara, et al., Improvement of the *Oryza sativa* Nipponbare reference genome using next generation sequence and optical map data. Rice. 6, 4 (2013).

25. E. F. Ullman, et al., Luminescent oxygen channeling assay (LOCI): sensitive, broadly applicable homogeneous immunoassay method. Clin. chem. 42, 1518–1526 (1996).

26. K. Nemoto, et al., Tyrosine phosphorylation of the GARU E3 ubiquitin ligase promotes gibberellin signalling by preventing GID1 degradation. Nat. commun. 8, 1004 (2017).

27. J. H. R. Maidment, et al., Multiple variants of the fungal effector AVR-Pik bind the HMA domain of the rice protein OsHIPP19, providing a foundation to engineer plant defense. J. Biol. Chem. 296, 100371 (2021).

28. M. Mirdita, et al., ColabFold: making protein folding accessible to all. Nat. methods 19, 679– 682 (2022)

29. C. Thomas, et al., Specific targeting of a plasmodesmal protein affecting cell-to-cell communication. PLoS Biol. 6, e7 (2008).

30. P. C Bull, D. W. Cox, Wilson disease and Menkes disease: new handles on heavy-metal transport. Trends Genet. 10, 246–252 (1994).

31. C. Askwith, et al., The FET3 gene of S. cerevisiae encodes a multicopper oxidase required for ferrous iron uptake. Cell. 76, 403–410 (1994).

32. R. A. Pufahl, et al., Metal ion chaperone function of the soluble Cu(I) receptor Atx1. Science. 278, 853–856 (1997).

33. A. C. Rosenzweig, T. V. O’Halloran, Structure and chemistry of the copper chaperone proteins. Curr. Opin. Chem. Biol. 4, 140–147 (2000).

34. P. E. Dykema, et al., A new class of proteins capable of binding transition metals. Plant Mol. Biol. 41, 139–150 (1999).

35. O. Barth, et al., Stress induced and nuclear localized HIPP26 from Arabidopsis thaliana interacts via its heavy metal associated domain with the drought stress related zinc finger transcription factor ATHB29. Plant Mol. Biol. 69, 213–226 (2009).

36. E. Himelblau, et al., Identification of a functional homolog of the yeast copper homeostasis gene *ATX1* from Arabidopsis. Plant Physiol. 117, 1227–1234 (1998).

37. S. Puig, et al., Higher plants possess two different types of ATX1-like copper chaperones. Biochem. Biophys. Res. Commun. 354, 385–390 (2007).

38. L.J. Shin, J.C. Lo, K.C. Yeh, Copper chaperone antioxidant protein 1 is essential for copper homeostasis. Plant Physiol. 159, 1099–1110 (2012).

39. W. Gao, et al., *Arabidopsis thaliana* acyl-CoA-binding protein ACBP2 interacts with heavy-metal-binding farnesylated protein AtFP6. New Phytol. 181, 89–102 (2009).

40. Y. Zhu, L. Liu, L. Shen, H. Yu, NaKR1 regulates long-distance movement of FLOWERING LOCUS T in *Arabidopsis*. Nat. Plants. 2, 16075 (2016).

41. G. H. Cowan, et al., Potato Mop-Top Virus Co-Opts the Stress Sensor HIPP26 for Long-Distance Movement. Plant Physiol. 176, 2052–2070 (2018).

42. X. Zhang, et al., Isolation and characterisation of cDNA encoding a wheat heavy metal-associated isoprenylated protein involved in stress responses. Plant Biol. 17, 1176–1186 (2015).

43. Q. Imran, et al., Nitric Oxide Responsive Heavy Metal-Associated Gene *AtHMAD1* Contributes to Development and Disease Resistance in *Arabidopsis thaliana*. Front. Plant Sci. 7, 1712 (2016).

44. Z. Rodakovic, et al., Arabidopsis *HIPP27* is a host susceptibility gene for the beet cyst nematode *Heterodera shachtii*. Molecular Plant Pathol. 19, 1917–1928 (2018).

45. C. Le Roux, et al., A receptor pair with an integrated decoy converts pathogen disabling of transcription factors to immunity. Cell. 161, 1074–1088 (2015).

46. P. F. Sarris, et al., A plant immune receptor detects pathogen effectors that target WRKY transcription factors. Cell. 161, 1089–1100 (2015).

47. E. Grund, D. Tremousaygue, L. Deslandes, Plant NLRs with integrated domains: Unity makes strength. Plant Physiol. 179, 1227–1235 (2020).

48. A. Longya, et al., Gene Duplication and Mutation in the Emergence of a Novel Aggressive Allele of the AVR-Pik Effector in the Rice Blast Fungus. Molecular plant-microbe interactions, 32, 740–749 (2019).

49. K. Katoh, D. M. Standley, MAFFT multiple sequence alignment software version 7: improvements in performance and usability. Mol. Biol. Evol. 30, 772–780 (2013).

50. L. T. Nguyen, H. A. Schmidt, A. von Haeseler, B. Q. Minh, IQ-TREE: a fast and effective stochastic algorithm for estimating maximum-likelihood phylogenies. Mol. Biol. Evol. 32, 268– 274 (2015).

51. D. T. Hoang, O. Chernomor, A. von Haeseler, B. Q. Minh, L. S. Vinh, UFBoot2: Improving the Ultrafast Bootstrap Approximation. Mol. Biol. Evol. 35, 518–522 (2018).

52. S. Kalyaanamoorthy, et al., ModelFinder: fast model selection for accurate phylogenetic estimates. Na Methods 14, 587–589 (2017).

53. D. Kim, B. Langmead S. L. Salzberg, HISAT: a fast spliced aligner with low memory requirements. Nat. methods. 12, 357–360 (2015).

54. M. Pertea, et al., StringTie enables improved reconstruction of a transcriptome from RNA-seq reads. Nat. Biotechnol. 33, 290–295 (2015).

55. C. Engler, R. Kandzia, S. Marillonnet, A one pot, one step, precision cloning method with high throughput capability. PloS One. 3, e3647 (2008).

56. T. Sawasaki, T. Ogasawara, R. Morishita, Y. Endo, A cell-free protein synthesis system for high-throughput proteomics. Proc. Natl. Acad. Sci. U. S. A. 99, 14652–14657. (2002)

57. K. Takai, T. Sawasaki, Y. Endo, Practical cell-free protein synthesis system using purified wheat embryos. Nat. protocols. 5, 227–238 (2010).

58. T. Sawasaki, et al., Arabidopsis HY5 protein functions as a DNA-binding tag for purification and functional immobilization of proteins on agarose/DNA microplate. FEBS lett. 582, 221–228 (2008).

59. R. Yin, B. Y. Feng, A. Varshney, B. G. Pierce, Benchmarking AlphaFold for protein complex modeling reveals accuracy determinants. Protein Sci. 31, e4379 (2022).

60. M. Mikami, S. Toki, M. Endo, M. Comparison of CRISPR/Cas9 expression constructs for efficient targeted mutagenesis in rice. Plant Mol. Biol. 88, 561–572 (2015).

61. S. Toki, et al, Early infection of scutellum tissue with Agrobacterium allows high-speed transformation of rice. Plant J. 47, 969–976 (2006).

62. C. A. Schneider, Image to ImageJ: 25 years of image analysis. Nat. methods. 9, 671–675 (2012).

## SI References

1. H. Kanzaki, et al., Arms race co-evolution of Magnaporthe oryzae AVR-Pik and rice Pik genes driven by their physical interactions. Plant J. 72, 894–907 (2012).

2. R. Yin, B. Y. Feng, A. Varshney, B. G. Pierce, Benchmarking AlphaFold for protein complex modeling reveals accuracy determinants. Protein Sci. 31, e4379 (2022).

